# Mapping 3D cellular mechanical activity with matrix-embedded DNA force-history probes

**DOI:** 10.64898/2026.08.05.742976

**Authors:** Yu-Hsuan Peng, Mathilde Lettinga, Anna Taubenberger, Elisha Krieg

## Abstract

Cells continuously integrate mechanical cues from the surrounding extracellular matrix to control fundamental biological processes such as proliferation and migration. Yet, studying the mechanical activity of cells in 3D over time is challenging, low-throughput, and requires specialized equipment. Here, we introduce DNA-based force-history probes, which convert transient pico-Newton forces into cumulative fluorescent signals within a mechanically adjustable DNA-crosslinked cell culture matrix. The probes provide control over signal lifetimes, allowing stress patterns to be recorded over minutes to days with tunable temporal memory. We use this system to visualize the mechanical activity of breast cancer spheroids and map the trajectories of migrating cancer cells. By combining different fluorophores and DNA-encoded signal lifetimes, we produce dual-color probes that associate temporal information to mechanical events. Overall, force-history probes provide an endpoint-readable record of mechanical cell-matrix activity, offering new opportunities for studying cell function in physiology and disease.

## Introduction

Cells actively probe and remodel their microenvironment by applying mechanical force to the extracellular matrix (ECM). These pushing and pulling movements generate local 3D stress fields, which trigger mechano-signaling feedback loops and guide cell proliferation, migration, and differentiation^1,2^. Increasing evidence shows that these interactions in a 3D matrix environment differ fundamentally from those on classical 2D cell culture substrates^3,4^. In 3D, mechanical confinement by the ECM restricts shape changes, while allowing cells to exert contractile, protrusive and volumetric forces^3^. The active remodeling of the ECM, in turn, alters patterns of force transmission and ultimately influences cell behavior^5,6^. Understanding this dynamic reciprocity between cells and surrounding matrices requires tools capable of mapping mechanical forces with spatial and temporal information.

Over the past decades, several biophysical tools have been developed to measure cell-exerted forces in 3D. Classical approaches such as 3D traction force microscopy quantify cell-generated forces by tracking the displacement of matrix-embedded particles^7–9^. These methods typically assume an isotropic and elastic matrix to enable the calculation of forces with nano-Newton sensitivity. However, molecular cell-matrix interactions involve pico-Newton (pN) forces, and a realistic cellular microenvironment is anisotropic and viscoelastic^10,11^. This discrepancy complicates force reconstruction in physiologically relevant materials. Different other innovative approaches have emerged to interrogate forces within multicellular structures using embedded pressure sensors or post hoc structural markers. Campàs et al. embedded microdroplets within living embryonic tissue to infer local stresses from droplet deformation^12^. Dolega et al. encapsulated compressible microbeads within spheroids to quantify the propagation of osmotic pressure^13^. Arnoldini et al. stained stretched fibronectin fibers with dye-labeled binding peptides to visualize stresses in histological tissue sections^14^. These approaches provide strain information either locally around specific probes (oil droplets or elastic beads), or as endpoint readouts in fixed tissue (dye-labeled peptides). However, comprehensive mapping of the mechanical activity of living cells across broad spatial and temporal scales remains an open challenge.

Recently, molecular force probes have emerged as a versatile tool to convert mechanical forces into fluorescence readouts^15–20^. These probes employ a *mechanoswitch*, often based on DNA hairpins or peptides that reversibly unfold or extend when the tension exceeds a characteristic threshold. The switch typically produces a signal through Förster resonance energy transfer. DNA-based force probes are particularly appealing, as the predictable Watson-Crick base pairing enables modular construction of mechanoswitches with rational control over their activation threshold^21^. Importantly, the force required to rupture DNA duplexes can be tuned between 1 and 60 pN^22–25^, coinciding with the forces typically exerted by cell surface receptors^21,24,26,27^.

Despite these promising developments, translation of fluorescent force probes from 2D culture dishes to more realistic 3D environments is difficult^20^. Previously reported probes measuring cell-ECM interactions required near-surface imaging, where scattering and background fluorescence are low enough for the weak, transient probe signals to be detected. Merindol and coworkers successfully used DNA hairpin probes to localize stresses in a hydrogel.^28^ This approach is promising, however, large deformations are required to activate a sufficient number of probes. Moreover, as the force relaxes, the probes re-fold or assemble back into their dark state, leading to premature loss of signal. Consequently, microscopic mechanical events (e.g., rapid protrusions of migrating cells or the fluctuating stresses during morphogenesis) would go undetected. To faithfully capture transient events, a cell-compatible and force-sensitive material is needed that can detect and memorize brief cell-matrix interactions and generate a cumulative signal.

Here, we describe force-history probes (FHPs), which can record the location and recency of mechanical deformations across an entire 3D cell culture volume. The probes self-integrate into DyNAtrix^29^, a viscoelastic DNA-crosslinked cell culture matrix with programmable mechanical properties. FHPs leverage a novel capping mechanism that allows accumulation of a strong signal and the dynamic adjustment of the fluorescent signal decay from minutes to days. We demonstrate that FHPs effectively visualize the distortional forces generated by growing breast cancer spheroids, mark the contractile and protrusive forces from cell spreading, and record the migration tracks of invasive cancer cells. This platform provides a robust and programmable tool to decipher the mechanical history of cell-matrix interactions.

## Results

### Material design

The force-sensitive matrix is formed by crosslinking DNA-functionalized polymers^30,31^ (**P**) with combinatorial^29,32^ dual-splint DNA crosslinkers (CLs) (50–80 µM) and FHPs (1 µM) (Figure 1, Supplementary Figure 1). DNA-based anchor strands on the polymer backbone serve as universal binding sites for the CLs and FHPs. Additional linear RGD peptide moieties on the backbone provide adhesion sites for integrins by which embedded cells apply force to the ECM^33^. The CLs were adjusted^6,29^ to set the shear storage modulus to 135 Pa and the stress-relaxation time to ∼1.1 days, providing a controlled environment for mechanobiological studies (Supplementary Figure 3).

**Figure 1.**
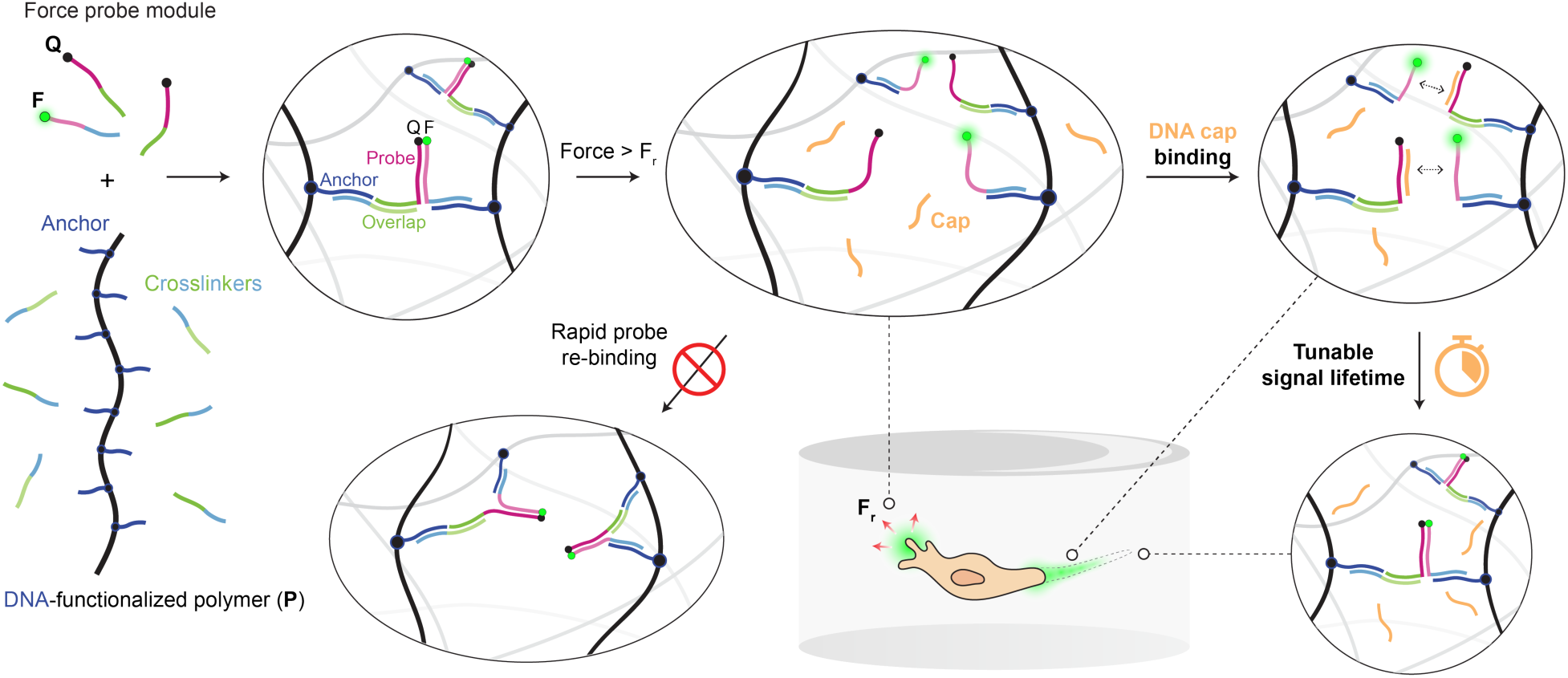
Working principle of force-history probes in a 3D cell culture matrix. FHPs and CLs hybridize to DNA anchor strands conjugated to a DNA-functionalized polymer to form a crosslinked network. Embedded cells apply localized forces to the ECM, for instance during cell migration. When the local force exceeds the rupture force of the probe (F_r_), it unzips, separating the fluorophore and the quencher, and exposing a cryptic binding site for DNA capping strands (“caps”). The **F** and **Q** strands can re-hybridize, either with their original binding partners, or with other activated probe strands in their vicinity. Freely diffusing caps bind to the activated probe strands, thereby blocking their recombination and suppressing premature signal decay. The fluorescence signal is retained for an adjustable period of time, controlled by the capping strand length and sequence. The decay time of the FHP determines for how long mechanical cell-matrix interactions are recorded in the matrix.

The FHP contains three essential components: fluorophore-modified strands (**F**), quencher-modified strands (**Q**), and capping strands (“caps”) (Figure 1). Hybridization of **F** to **Q** forms the mechanoswitch, whose duplex length and sequence determine its mechanical rupture force. The current FHP design is based on a 26-nt sequence with a GC content of 35%. It is predicted to unzip at a threshold of approximately 3.3 pN at 37 °C, which is substantially lower than the expected rupture force of the CL’s 22-nt overlap domain (26 pN) and of the anchor strand linkage to the polymer backbone (30 pN) (Supplementary Methods 2.12, Supplementary Figure 4)^34^. Representing the weakest link in the gel network, the FHPs were expected to detect cell-matrix interactions approaching and exceeding the 3.3 pN threshold. Early experiments revealed that the FHP-functionalized matrix indeed produces a fluorescent signal after mechanical breakage; however, activated probes quickly re-hybridized, causing rapid loss of the already weak signal. To solve this problem, we introduced caps, which are designed as freely diffusing DNA-based surrogates that quickly bind activated probes to prevent their premature rehybridization. Through this mechanism, caps convert the transient mechanoswitch activation into a cumulative fluorescence signal with adjustable decay kinetics.

### Force-history probes provide broad control over signal lifetime with low background

To evaluate the performance of FHPs, the hydrogels were perfused with caps, and the fluorescence signal was recorded over 24 hours, with and without prior mechanical activation. For mechanical activation, the gels were subjected to repeated pipetting with a positive displacement pipettor, yielding a strong fluorescence signal in all tested samples (Figure 2a, d; Supplementary Video 1, Supplementary Video 2). In the absence of capping strands, the initial force-activated signal was weaker, and it decayed to baseline levels within hours (Figure 2d).

**Figure 2.**
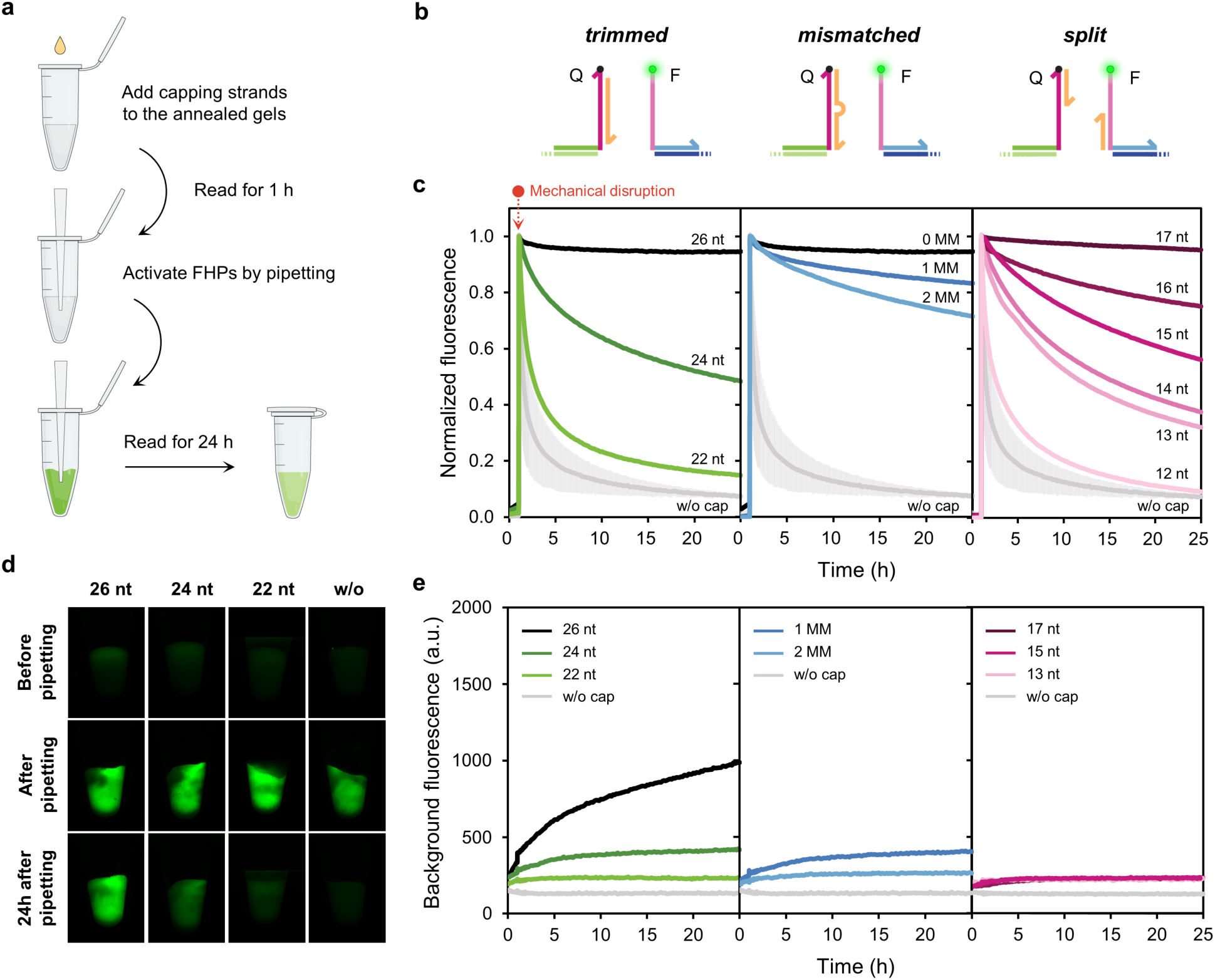
Capping strands control the signal lifetime of activated force probes. **a)** Scheme of the mechanical disruption experiment. Capping strands were added to the self-assembled hydrogels and the material’s fluorescence was recorded for one hour. The material was then mechanically disrupted by positive-displacement pipetting, and the fluorescence recording was continued for another 24 hours. **b)** Scheme of the three capping strategies, and **c)** normalized fluorescence decay profiles in the presence of *trimmed* (left), *mismatched* (center), and *split* (right) capping strands. **d)** Representative fluorescence images of hydrogels in PCR tubes exhibiting adjustable signal decay with 22nt–26nt trimmed capping strands. **e)** Increase of the background fluorescence in the hydrogels due to leakage. The *split* cap design minimizes leakage compared to trimmed and mismatched designs. All experiments were conducted at 20°C.

We explored three alternative capping strand designs, termed *trimmed*, *mismatched*, and *split* capping (Figure 2b). *Trimmed* caps were designed to bind **Q** via 22 to 26 base pairs. Longer caps were expected to form more stable cap/**Q** complexes, thereby retarding the displacement reaction by the rehybridizing probe strands. The *mismatched* approach relies on a similar concept, but the cap is kept at a fixed length (26 nt), and the cap/**Q** complex stability is adjusted by introducing base pair mismatches. Finally, s*plit* caps were designed to bind different regions of **F** and **Q**, and the complex stability was controlled via duplex length from 12 to 17 base pairs. All three tested approaches provided the anticipated level of control over force-activated signal lifetimes. Changing cap length allowed the tuning of the signal half-time from 1.5 h to far beyond 24 h (Figure 2c). The *mismatched* approach showed a similar trend, though the tested number of mismatches did not give access to sub-24 h signal half-times.

Control measurements with mechanically unperturbed gels revealed that *trimmed* and *mismatched* caps caused substantial increase in background fluorescence (Figure 2e). Increasing the length and decreasing mismatches therefore did not only alter the probe’s re-hybridization kinetics, but also biased its assembly thermodynamics towards the open state. This undesired activation in the absence of mechanical force is a form of leakage^35^. It is possibly due to DNA ‘fraying’ and ‘breathing’—dynamic processes that make paired nucleobases available for spurious cap binding and subsequent strand displacement (Supplementary Figure 5a). Leakage did not affect the *split* capping approach to the same extent. Though spurious binding of the *split* caps to the probe is possible, their reduced length does not readily cause full dissociation of the probe (Supplementary Figure 5b). The *split* design thus caused minimal background fluorescence, while maintaining the ability to tune the retention of the force-activated signal over a broad range of timescales (Figure 2e). S*plit* capping was consequently used in all later experiments.

### Force-history probes track cell-matrix interactions

To test the performance of FHPs in cell culture, we embedded breast cancer spheroids derived from MCF-10A ER-Src cells in DyNAtrix. DyNAtrix was crosslinked via heat-activated CLs^6,29^ and FHPs (Supplementary Figure 6). Tamoxifen was added to the culture medium for Src induction, driving transition from a non-invasive towards an invasive phenotype within 48 hours^36,37^ (Supplementary Figure 7). The matrix was also supplemented with a control probe, nearly identical to the FHP but attached to the network as a dangling end, which prevents its activation by mechanical load (Figure 3a, Supplementary Figure 8). The control probe served as an internal reference to monitor fluorescence changes arising from non-mechanical events, such as fluctuations in matrix concentration, probe degradation, or leakage. Signal lifetime tuning was readily achieved under cell culture conditions (DMEM, 37 °C) but required the use of slightly longer (16–19nt) cap strands to enable broad coverage of signal lifetimes at the elevated temperature (Supplementary Figure 9).

**Figure 3.**
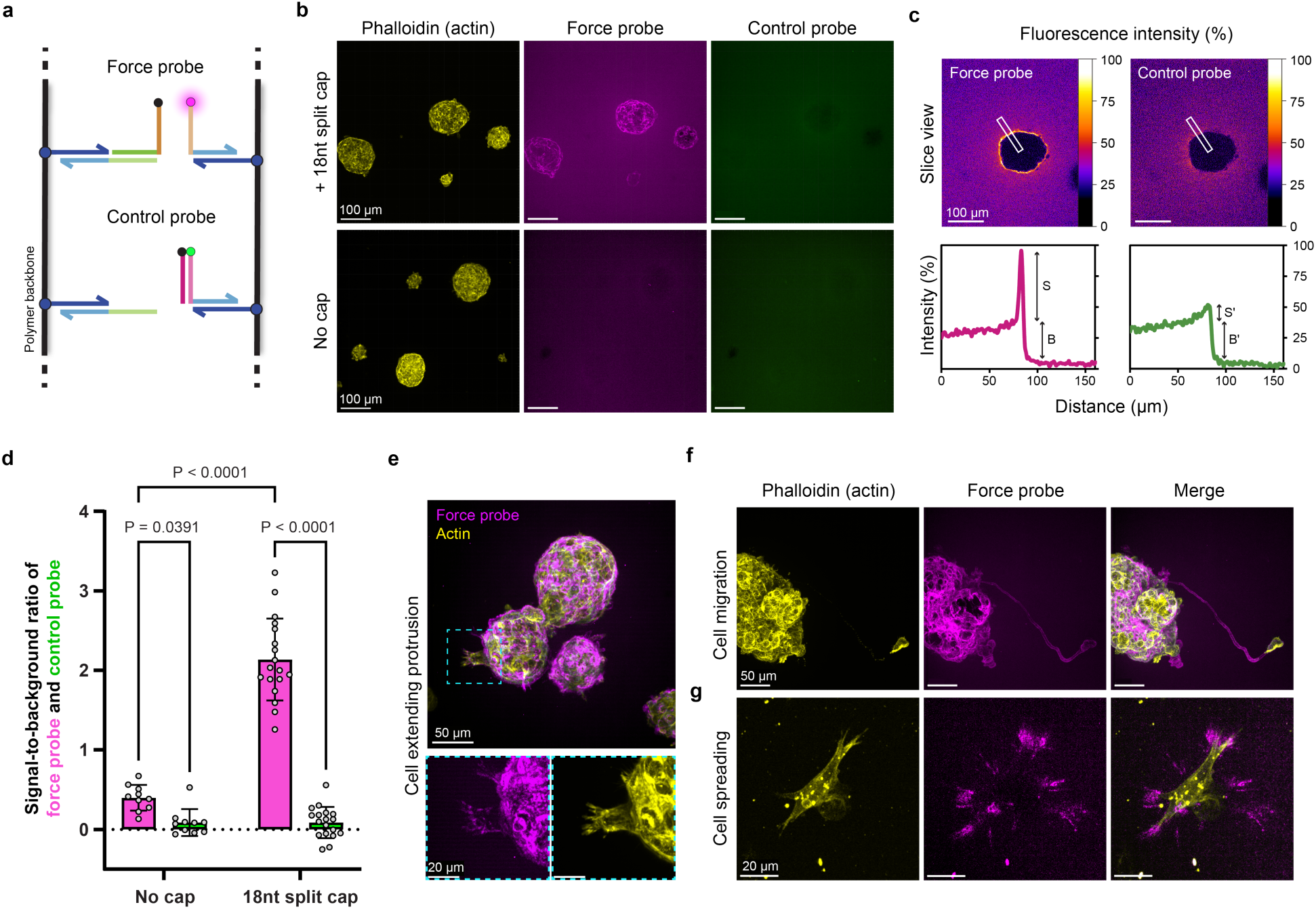
Recording of mechanical cellular activity in 3D culture with force-history probes. **a)** Scheme of the hydrogel setup containing FHP and force-insensitive control probes. Caps are omitted for clarity. **b)** Confocal images of breast cancer spheroids cultured in DyNAtrix for 6 days. Cells were stained with phalloidin (F-actin). **c)** Cross-sectional analysis of the fluorescence intensity of FHP and control probe across the spheroid-matrix interface. Top: heatmap of fluorescence intensity on a confocal section. Bottom: intensity plots across the ROI (indicated with a white box). **d)** Signal-to-background ratio of FHP and control probe at spheroid-matrix interfaces with and without 18 nt caps. Data are shown as mean ± s.d. (no cap: n = 10; 18nt split cap: n = 19 spheroids). Statistical analysis was performed using a one-way ANOVA with Fisher’s LSD test. **e)** Top: confocal image of MCF-10A ER Src breast cancer spheroids. The cyan box highlights a cell extending an F-actin rich protrusion into the matrix. Bottom: enlarged view of fluorescence pattern at the protrusion arm. **f)** Representative confocal image of a MCF-10A ER Src cancer spheroid and a migrating cell leaving a fluorescence trace in gel. **g)** Spreading breast cancer cell whose contractile forces activate FHPs, producing a 3D stress pattern that extends up to 10–20 µm into the surrounding matrix. All confocal images are shown as maximum projections.

We were initially concerned about potential degradation of FHPs, facilitated either through cellular uptake or secretion of nucleases. Early culture experiments showed that a Cy5-labeled FHP was indeed prone to severe cellular uptake, likely due to the affinity of this fluorophore to cell membranes and certain organelles^38^. The use of non-lipophilic and negatively charged fluorophores like FAM, however, effectively prevented probe uptake (Supplementary Figure 10). Notably, force probes were substantially degraded when exposed to serum-supplemented medium. We had previously shown that DNA can be protected against nucleases in cell culture media by addition of actin^29^. However, this protective measure was not required, since DNA probes were neither degraded in fresh serum-free medium, nor in used medium that was analyzed after 48 hours of cell culture, demonstrating that nuclease secretion by breast cancer cells was minimal (Supplementary Figure 11).

Confocal microscopy demonstrated that FHPs sensitively respond to the mechanical activity of tumor spheroids in DyNAtrix (Figure 3b-f). The signal is created as individual cells from the spheroid push into the matrix, leaving behind increasing numbers of fluorescent imprints (Supplementary Video 3). Signal-to-background (S/B) ratios at the spheroid-matrix interface were computed based on the fluorescent channels of the force probe and the control probe (Figure 3c; Supplementary Figure 12). Control probes produced faint but non-zero S/B ratios, likely due to the compaction of the matrix around the growing spheroids. Without caps, force probes also produced weak signals, albeit significantly stronger than the control probes (p=0.039; Figure 3d). In contrast, the presence of caps yielded significantly enhanced and durable force-activated signals without affecting the control channel (p<0.001). Consistent with the prior validation of signal lifetimes, increasing the lengths of caps improved the FHP’s performance, providing optimal S/B ratios at 18 nt length (Supplementary Figure 13). Though longer caps yield longer signal lifetimes, nearest-neighbor thermodynamic calculations show that beyond 18 nt, caps become prone to dimerize (Supplementary Figure 14). The dimerization is expected to impair their ability to rapidly block activated probes, explaining the slight decrease of S/B ratio for 19 nt caps.

Besides blunt expansion forces, FHPs also revealed exceptionally fine mechanical interaction features. High-magnification images showed cells in the periphery of the spheroid extending F-actin rich protrusions into the matrix, leaving filament-like pattern of activation signal with diameters of 1.2–1.5 µm (Figure 3e). The force pattern matches the typical rim tension generated by filopodia and lamellipodia structure on the leading edge of protrusion arms. FHP activation also marked contractile stress fields arising from cell spreading, propagating up to 20 µm into the matrix (Figure 3g). After protruding into the matrix, some individual cells fully detached from the spheroid and invaded deeply into the gel, creating fluorescent tracks that marked their migration trajectories (Figure 3f, Supplementary Figure 15). The emerging migration channels displayed relatively uniform diameters of 6.02 ± 0.26 µm (Supplementary Figure 16). Some of the FHP-marked migration tracks contained numerous actin-rich puncta. These structures are consistent with retraction fibers, cellular fragments that are shed from migrating cells, which have been previously reported to guide cellular migration of cancer cells on 2D surfaces^39^. Notably, in several instances a subsequent breast cancer cell was observed to deform, squeeze into, and follow the fluorescent tracks of an earlier migrating cell (Supplementary Figure 15). This behavior is consistent with the idea that a pioneer cell can create a persistent, mechanically permissive path that lowers the physical barrier for later cells. Previous studies showed that cancer cells can exploit matrix microtracks, including tracks generated by proteolytic remodeling by other cancer cells^40^ or by mechanically active cancer-associated fibroblasts^41^. Our observations provide direct evidence that breast cancer cells themselves can generate persistent guidance tracks in a fully synthetic, proteolytically non-degradable matrix through mechanical matrix deformation.

Finally, we demonstrated the broad applicability of the DyNAtrix platform across cell types by utilizing FHPs to visualize mechanical signatures of two additional, highly invasive yet biologically distinct cancer models: MDA-MB-231 human breast cancer cells and U87MG human glioblastoma cells (Supplementary Figure 17; Supplementary Videos 4 and 5). MDA-MB-231 cells exhibited an elongated morphology, generating pronounced probe signals via traction forces along their leading and trailing edges. In contrast, glioblastoma cells extensively probed the microenvironment using their characteristic ultra-long, thin cellular protrusions, leaving spindle-like traces of probe activation.

### Two-color force-history probes allow temporal encoding of mechanical events

Finally, we wished to test whether FHPs could be used not only to map the spatial distribution of mechanical events, but also their sequence in time. To this end, we developed a dual-color FHP system, comprising two orthogonal probe sequences terminated by distinguishable fluorophores, Alexa Fluor 647 and FAM. We employed two sets of caps complementary to the two FHPs. The first probe was combined with shorter caps, producing a short-lived red signal. The second probe was combined with longer caps, producing a long-lived green signal. We envisioned that the relative signals from the short- and long-lived probes would provide a coarse indication of when a mechanical disruption had occurred, distinguishing recent from earlier events (Figure 4a).

**Figure 4.**
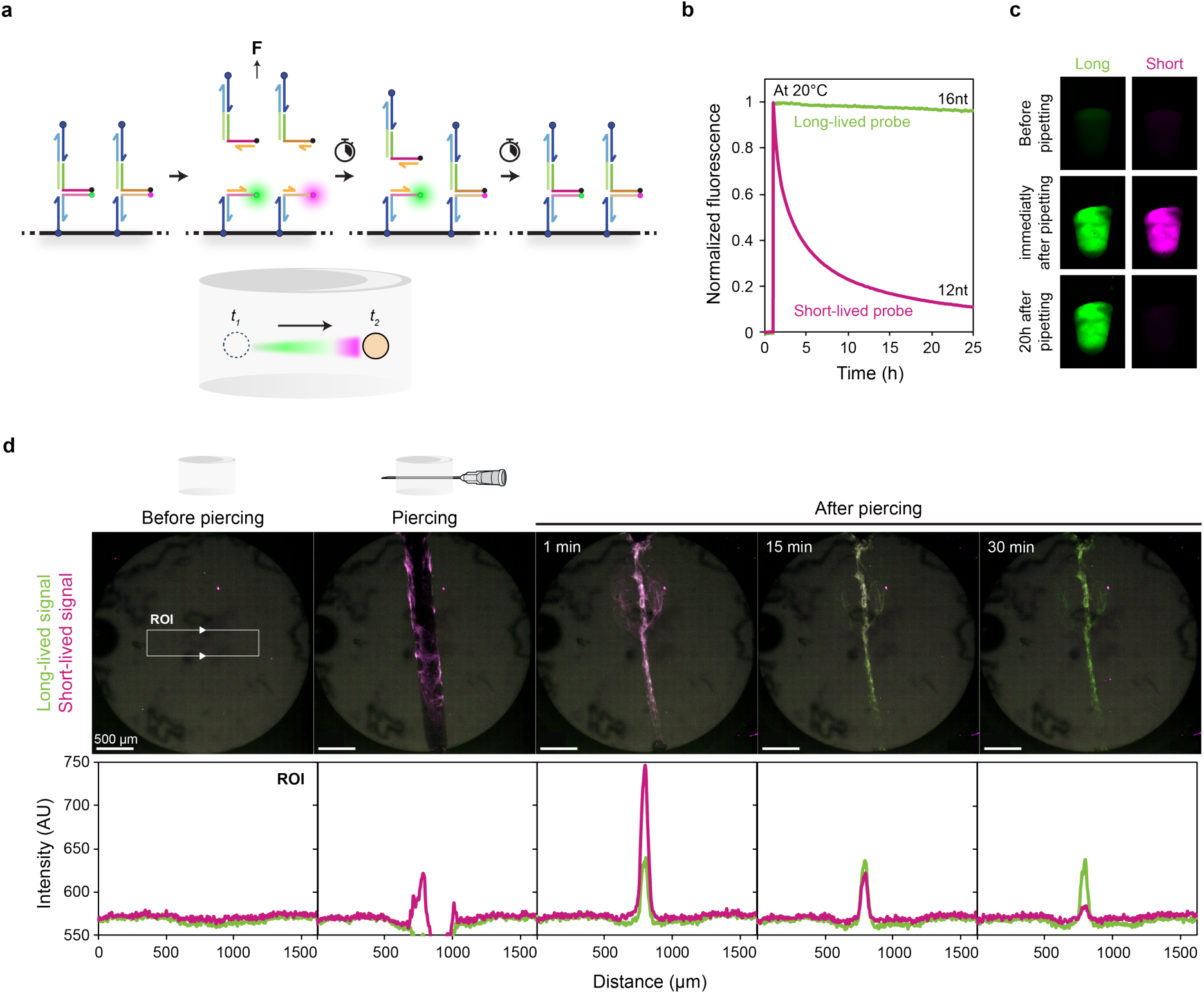
Dual-color force-history probes encode the recency of mechanical events. **a)** Scheme of DyNAtrix combining a fast-decay FHP and a slow-decay FHP. The intensity of the two signals could encode the sequence of FHP activation, for instance, due to cell migration. **b)** Normalized fluorescence intensity of a slow-decay FHP (16nt caps; FAM) versus a fast-decay FHP (12nt caps; Alexa Fluor 647) after mechanical disruption by positive-displacement pipetting at 20°C. **c)** Fluorescent images of the corresponding bulk hydrogels in a PCR tube before and after pipetting. **d)** Piercing test. Top: Confocal microscopy images of a gel sample before, during, and after being punctured by a syringe needle at 25°C. Bottom: Raw intensity profiles along the cross-section of the damaged site (region of interest (ROI) indicated by a white box).

To validate the dual-color FHP system, we first conducted mechanical disruption experiments at room temperature. 16-nt and 12-nt caps were used to confer slow and fast signal decay, respectively. Mechanical activation led to strong initial emission from both FHPs with a distinctive fluorescence decay (Figure 4b,c). We next conducted a piercing experiment, where the hydrogel was imaged in the confocal microscope before, during, and after being punctured by a needle (Figure 4d). Immediately after the puncture, the damaged site showed strong fluorescence in both channels. The short-lived FHP signal decayed to near-background values within the subsequent 30 minutes, whereas the green signal remained stable. This proof-of-concept demonstrates that it is possible to design multiple FHPs that are independently addressable, without interfering with each other’s function. The distinct color-coded signal lifetimes provide a built-in timestamp that can allow users to distinguish mechanical events in the distant past from more recent events in a single snapshot.

## Discussion

In this work, we introduced force history probes (FHPs), a tool for 3D mapping of mechanical cell– matrix interactions. By combining DNA force probes and capping strands in DyNAtrix, transient force-induced probe activation is converted into a cumulative signal. The nucleotide sequence of the DNA probe encodes its activation threshold^21^, whereas the capping strand length encodes the characteristic signal decay time. FHPs enable spatial visualization of time-integrated force-threshold events across timescales ranging from minutes to days, providing a record of cellular activity that is inaccessible to conventional probes. We demonstrated force-history measurements in cell culture for up to six days, with readout either after sample fixation or during live-cell imaging.

DNA-based tension probes have become a powerful technology for mechanobiological studies in 2D cell culture^15,16,18–21^. Salaita and coworkers previously used locking oligonucleotides to integrate forces in 2D cell culture experiments, demonstrating the value of stabilizing otherwise transient probe activation^42,43^. Our hairpin-free split capping approach extends this capability, enabling the advancement of DNA-based force probes into 3D matrices for organoid research and disease models^44^. The dual-color FHP system with different signal lifetimes can be useful for distinguishing long-term cellular activities (e.g., cell migration over multiple days) from more recent activities (e.g., cell protrusion and contraction).

In purely elastic hydrogels with mesh sizes smaller than the nucleus (∼3 µm), cells cannot migrate without proteolytic degradation^45^. In contrast, native breast tumor tissue can undergo plastic deformation, permitting migration through both proteolytic and protease-independent mechanical mechanisms^46^. DyNAtrix likewise undergoes plastic deformation, but its nanoporous structure, with a mesh size of approximately 32 nm (Supplementary Note 1), is proteolytically non-degradable. FHP-functionalized DyNAtrix can therefore help isolate and study degradation-independent, mechanically mediated cell–matrix interactions.

We identify the mechanical footprints of breast cancer cells, highlighting distortional stresses produced by spheroid expansion, as well as the protrusion and contraction exerted by individual cells actively migrating through the matrix. The ability to record migration trajectories yields mechanistic insight into how cells navigate their 3D environment. Moreover, FHPs can provide “mechanical fingerprints”, which are increasingly recognized as useful indicators of normal and diseased cellular states in pathological pathways, including cancer metastasis and fibrosis^2^. The combination of the adjustable viscoelastic properties of DyNAtrix with time-integrated force sensing may also complement conventional cell-invasion assays by linking invasive behavior to patterns of mechanical matrix engagement.

FHPs offer complementary strengths and limitations relative to existing deformation-based force-mapping methods^7–9,12–14^. In comparison to 3D traction force microscopy, FHPs enable highly *sensitive* and *direct* measurements, since probe activation takes place at a predictable^25,34^ pico-Newton force. This threshold-based readout can be applied in viscoelastic environments like DyNAtrix, which is adjustable to faithfully mimic the wide range of stress-relaxation properties of native tissues^6,29^. The high probe concentration (∼600 FHPs per µm^3^) enables dense volumetric sampling throughout the imaged matrix. As opposed to microdroplet-deformation probes, which report forces around discrete locations, FHPs map forces throughout the imaged volume, which makes them suitable for high-throughput experiments without manual interrogation. At the level of an individual probe, the FHP response is binary: it reports whether the activation threshold has been exceeded. At the ensemble level, signal intensity reflects the number and recency of activation events, but it does not directly quantify force magnitude, strain magnitude or vector direction. Because the signal is integrated over a finite memory window, it also cannot recover the precise timing of individual events, and similar signals may arise from sustained loading or repeated transient loading. FHPs thus provide spatiotemporal information that complements, rather than replaces, established deformation-based methods.

Although FHPs were designed to self-integrate into the DyNAtrix cell culture platform, the concept can be equally applied in other DNA-functionalized gels. FHP-infused DyNAtrix could be used to form interpenetrating networks with traditional cell culture systems like Matrigel, collagen, or GelMA, endowing these widely used platforms with mechanosensitive functionality. For applications that require long-term stability in the presence of serum, FHPs could be built from nuclease-resistant L-DNA analogs^47^. While the presented FHPs detect deformation forces across the entire 3D matrix, alternative implementations could link the probe to RGD peptides or other receptor-binding ligands, thereby isolating interactions that are transmitted via specific cell-surface receptors^48,49^. We anticipate that FHPs will become a versatile tool to elucidate cell– matrix mechanical interactions in developmental biology and disease modeling.

## Methods

### Materials

Solvents and reagents were purchased from commercial sources and used as received, unless otherwise specified. Water was obtained from a Milli-Q system from Merck Millipore. Molecular biology grade acrylamide (catalog number A9099) and 19:1 acrylamide/bis-acrylamide (catalog number A2917), sodium acrylate (catalog number 408220), and ammonium persulfate (APS; catalog number A3678) were purchased from Sigma-Aldrich. Methanol (ACS reagent grade; catalog number 423955000), ultrapure N,N,Nʹ,Nʹ-tetramethylethylenediamine (TEMED; catalog number 15524010) were purchased from Thermo Fisher Scientific. Desalted oligonucleotides were purchased from Integrated DNA Technologies (IDT). Nitrogen gas (>99.999%) was used under inert conditions and supplied by an in-house gas generator. To ensure an inert condition, nitrogen gas was purified through a Model 1000 oxygen trap from Sigma-Aldrich (catalog number Z290246). Reagents with unreacted acrylamide groups were stored at 4 °C or −20 °C, protected from unnecessary exposure to light.

### Cell sources

The MCF-10A ER-Src cell line was kindly provided by Kevin Struhl (Harvard Medical School, ref. ^50^; RRID:CVCL_N805). MDA-MB-231 cell line was purchased from DSMZ (Leibniz Institute, German collection of microorganisms and cell cultures GmbH). Glioblastoma U87MG_mcherry cells were provided by Prof. Dr. Leoni Kunz-Schughart (OncoRay—National Center for Radiation Research in Oncology, Dresden, Germany, ref. ^51^). The original U87MG cell line was purchased in 2010 (CLS Cell Lines Service).

### Polymer synthesis

We synthesized two derivatives of the polymer, **P_10_** and 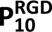, which differ in their degree of functionalization with anchor strands and cell-adhesion peptides, as previously described^29^. **P_10_** lacks cell-adhesion sites and was used for force-probe validations and optimizations that did not involve cells. 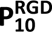 contains the arginine-glycine-aspartate (RGD) peptide motif to facilitate cell adhesion. All experiments involving cells used this polymer backbone. The synthesis protocol was adapted from refs. ^30^ and ^29^. In brief, acrylamide (50 mg ml^−1^), sodium acrylate (0.5 mg ml^−1^), and acrylamide-labeled anchor strand DNA (Supplementary Table 1, strand 1) were co-polymerized at a molar ratio of 10,000:100:10 in 1x TBE buffer (100 mM Tris, 90 mM boric acid, 1 mM EDTA, pH 8.3) to create DNA-grafted poly(acrylamide-co-acrylic acid). For the corresponding RGD-functionalized derivative 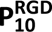, 10 mM acrylated RGD peptide (sequence: G(acryl-K)GGGRGDSP) was co-polymerized with the above solution. The synthesis of **P_10_** and 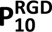 was initiated by addition of 0.025 wt% TEMED and 0.025 wt% APS. To achieve high molecular weight and a narrow size distribution, the reaction was performed in high-purity nitrogen gas, which was passed through an oxygen trap on-site. The reaction was allowed to proceed overnight and yielded a highly viscous solution, indicating the formation of long polymer chains. NMR spectroscopy was used to verify high conversion of the monomer (Supplementary Figure 2). The solution was diluted in 9 volumes of 1x TE buffer (10 mM Tris, 1 mM EDTA, pH 8.0) and subsequently purified via methanol precipitation. Finally, the pellet was resuspended in milliQ water at 2.5% (w/v) and stored in aliquots at −20 °C.

### Pipetting assay

Gel samples were prepared at a final concentration of 1% (w/v) **P_10_**, mixed with 50 µM of forward crosslinker (strand 2a), 60 µM reverse crosslinker (strand 2b), 1 µM fluorophore-labeled probe strand (F-strand 4a or 5a), and 2 µM quencher-labeled probe strand (Q-strand 4b or 5b), in 1x TE buffer and 150 mM NaCl. For two-color force-history system, 1 µM F-strands (both strands 4a and 5a) and 2 µM Q-strand (both strands 4b and 5b) were used. The samples were annealed on a C1000 Touch™ Thermal Cycler (Bio-Rad) using the following steps: (1) heating at 80 °C for 3 min, (2) cooling ramp from 80 °C to 20 °C at −1 °C min^−1^, (3) holding at 20 °C.

The corresponding DNA capping strands were added to the thermally annealed gel samples. Trimmed caps (strand 6, 7, or 8) and mismatched caps (strand 6, 9 or 10) were added to the gels to reach a final concentration at 4µM. For split caps, the Q-strand cap (strand 12a, 13a, 14a, 15a, 16a or 17a) was added to reach a final concentration at 4µM, while the F-strand cap (strand 12b, 13b, 14b, 15b, 16b or 17b) was added to reach a final concentration at 2µM. For the two-color force-history system, the Q-strand caps (strand 11a and 12a) and the F-strand caps (strand 11b and 12b) were added to reach a final concentration of 4 µM and 2 µM, respectively. Samples were read for 1 hour at 20 °C on a CFX96 Touch™ Real-Time PCR System (Bio-Rad). After 1 hour of reading, gels were mechanically perturbed by pipetting up and down for 20 times using a 10 µL positive displacement pipette (Gilson). Perturbed samples were then immediately read again at 20°C for an additional 24 hours. Data were processed using Bio-Rad CFX Maestro software (v4.1.2433.1219). Fluorescence images of PCR tubes (excitation at 473 nm/637nm) were acquired before and after perturbation as well as after 24h of recovery on a Typhoon FLA 9500 scanner (GE Healthcare Life Sciences) at a 50 µm pixel size with the accompanying software (v1.0).

*Data analysis*: the fluorescence intensities were normalized for comparison across different conditions. For each sample, the maximum value (I_max_) was defined as the peak fluorescence observed immediately after mechanical perturbation. The minimal value (I_min_) was defined as the mean fluorescence of the unperturbed control sample without capping strands. The fluorescence value at each timepoint (I) was rescaled according to: (I-I_min_)/(I_max_-I_min_). Signal half-life was defined by the time required for the normalized fluorescence to reach the midpoint of the decay curve (I_norm_ = 0.5).

### Preparation of DyNAtrix precursors for cell culture experiments

The DyNAtrix protocol was based on refs.^29^ and with adaptations for including FHPs. In brief, two precursor solutions, A and B, were prepared: precursor A was prepared by mixing 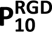 (final concentration: 1% (w/v)), forward crosslinker (strand 2a, final concentration: 80 µM), blocking strand (strand 3a, final concentration: 160 µM), quencher-labeled probe strand (strand 5a, final concentration: 1 µM), fluorophore-labeled probe strand (strand 5b, final concentration: 2 µM), quencher-labeled control strand (strand 4c, final concentration: 2 µM), and fluorophore-labeled control strand (strand 4b, final concentration: 1 µM). Precursor B was prepared by mixing reverse crosslinker (strand 2b, final concentration: 90 µM) and blocking strand (strand 3b, final concentration: 180 µM). 10x concentrated DMEM was added to both precursors to reach a final 1x concentration. The precursors were pre-annealed on a C1000 Touch™ Thermal Cycler (Bio-Rad) using the following steps: (1) heat to 70 °C for 3 min, (2) cool from 70 °C to 4 °C at −3 °C min^−1^, (3) hold at 4 °C. The precursor was stored at 4°C.

### Preparation of breast cancer spheroids

The MCF-10A ER-Src cells were maintained in T75 tissue culture flasks (BioLite Thermo Scientific) in serum-supplemented medium (Dulbecco’s Modified Eagle Medium/Nutrient Mixture F-12 (DMEM/F-12) (Gibco, Cat. #11039047) supplemented with 5% charcoal stripped horse serum (CSHS) (Invitrogen, Cat. #16050122), 1x Penicillin and Streptomycin (Gibco, Cat. #15140122), 20 ng/mL EGF (Miltenyi, Cat. #130093825), 0.5 µg/mL hydrocortisone (Sigma-Aldrich, Cat. #H0888), 100 ng/mL Cholera toxin (Sigma-Aldrich, Cat. #C8052), 10 µg/mL insulin (Sigma Cat. #I2643) and 0.5 µg/mL puromycin (Sigma-Aldrich, Cat. #540411)) at 37 °C, 5% CO_2_ until 70–80 % confluence. To generate spheroids, cells were detached in 0.25% trypsin-EDTA solution for 3 min, washed with PBS and mixed with 5% (w/v) gelatin methacryloyl (GelMA) (prepared according to ref. ^52^) and 0.15% (w/v) photoinitiator lithium phenyl-2,4,6-trimethylbenzoylphosphinate (LAP) (Sigma-Aldrich, Cat. #900889) at a density of 4 x 10^5^ cells/mL. Gels were formed by exposure to light of 405 nm wavelength for 90 seconds using the Luna photocrosslinker (Gelomics). Cells were cultured at 37 °C, 5% CO_2_ for 4 days. Formed spheroids were released from GelMA using 2.5 mg/mL collagenase solution (Nordmark Chemicals, Cat. #S1746501), washed twice with serum-free DMEM/F-12 and embedded into DyNAtrix (see below).

### 3D cell culture in force-sensitive DyNAtrix

5 µL of the above obtained MCF-10A ER-Src spheroid suspension was first mixed with 15 µL precursor A on ice. 5µL precursor B was then added to the mixture (total volume: 25 µL) and pipetted up and down for 25 times on ice. The final spheroid density was 1 x 10^5^ spheroids/mL in DyNAtrix. The mixture was then subjected to a micro-well insert (ibidi GmbH, catalog number 80409) placed in a 6-well plate with 5 μL cell-laden mixture per micro-well. To trigger heat-activated gelation, the plate was incubated at 37 °C with 5% CO_2_ for 60 min. Following gelation, 5 mL of the serum-free media (1x DMEM/F-12, 1x Insulin-Transferin-Selenium (Gibco, Cat. #41400045), 20 ng/mL EGF, 0.5 µg/mL hydrocortisone, 1x Penicillin and Streptomycin, 1 µM 4-hydroxytamoxifen (Sigma-Aldrich, Cat. #H7904)) was added to each well. While spheroids initially display a non-invasive and rounded morphology, they underwent morphological transformation upon Src induction via the supplementation of tamoxifen, leading to elongated cell shapes and enhanced cell motility^36,37^ (Supplementary Figure 7). After 24 hours of culture at 37 °C and 5% CO_2_, the corresponding DNA capping strands were added to the medium to reach a final concentration of 2 µM for F-strand caps (strand 16a, 17a, 18a or 19a), and 4 µM for Q-strand caps (strand 16b, 17b, 18b or 19b). The spheroids were cultured for 6 days and subsequently fixed for immunofluorescent staining.

### Immunofluorescent staining

MCF-10A ER-Src cells were fixed with 4% paraformaldehyde (PFA) for 30 min, washed with phosphate-buffered saline (PBS) buffer three times, and incubated in the permeabilization and blocking buffer (0.5% Triton™ X-100, 2% (w/v) bovine serum albumin (BSA) in PBS) for 30 min at room temperature. Then the cells were stained with Alexa Fluor 555 Phalloidin (Invitrogen, Cat. #A34055) at 1:200 dilution in 2% (w/v) BSA/PBS buffer overnight at 4 °C. Finally, the cells were washed with PBS three times. Confocal images were acquired on an Andor Dragonfly Confocal Microscope (Oxford Instruments) at 20x/40x magnification.

### Time-lapse confocal imaging

For live-cell imaging, MCF-10A ER-Src cells were stained with 1 µM CellTracker Orange (Thermo Fisher Scientific, Cat. #C2927) in serum-free medium for 30 minutes on day 2. The staining solution was then removed and replaced with fresh serum-free medium containing 18-nt *split* capping strands (2 µM strand 18a, and 4 µM strand 18b). Confocal images were acquired every 2 hours for 48 hours on an Andor Dragonfly Confocal Microscope (Oxford Instruments) at 37 °C, 5% CO_2_ at 20x magnification.

### Hydrogel piercing

The hydrogels were prepared at a final concentration of 1% (w/v) **P_10_**, mixed with 50 µM of forward crosslinker (strand 2a), 60 µM reverse crosslinker (strand 2b), 1 µM fluorophore-labeled probe strands (strand 4a and 5a), and 2 µM quencher-labeled probe strands (strand 4b and 5b), in 1x TE buffer and 150 mM NaCl. The hydrogels were thermally annealed in optical-bottom PCR tubes (Azenta, Cat. #4ti-0970) on a C1000 Touch™ Thermal Cycler (Bio-Rad) using the following steps: (1) heating at 80 °C for 3 min, (2) cooling ramp from 80 °C to 20 °C at −1 °C min^−1^, (3) holding at 20 °C.

After annealing, the Q-strand caps (strand 11a and 12a) and the F-strand caps (strand 11b and 12b) were dropped onto the gels to reach a final concentration of 4µM and 2µM, respectively. The samples were equilibrated at room temperature for an hour. Confocal images were acquired at room temperature (∼25 °C) on an Andor Dragonfly Confocal Microscope (Oxford Instruments) at 4x magnification. The samples were imaged before and after being pierced horizontally using a 30-gauge needle (Omnican).

### Statistics and reproducibility

Graphs and statistical analyses were made in GraphPad Prism, Excel, and Adobe Illustrator. Sample sizes are specified in the figure captions, where n refers to the number of distinct samples. For the S/B ratio analysis, a one-way ANOVA with Fisher’s Least Significant Difference (LSD) test was performed. Data are presented as mean ± s.d., with p-values denoted on the graphs. No statistical method was used to predetermine sample size, and no data was excluded from the analyses. Experiments were not randomized, and the investigators were not blinded to allocation during experiments or outcome assessment.

## Acknowledgements

E.K. acknowledges funding by the Federal Ministry of Research, Technology, and Space (BMFTR) in the program NanoMatFutur (grant no. 13XP5098) and from the Federal Ministry for Economic Affairs and Energy (BMWE) in the program EXIST Forschungstransfer (grant number 03EFSN0252). M.L. and A.T. are supported by DFG TA 751/7-1 (FOR5628) and the German Cancer Aid (Mildred Scheel Nachwuchszentrum Dresden P^2^). The authors thank Prof. Dr. Alf Honigmann, Prof. Dr. Carsten Werner, and Dr. Michele Marass for valuable discussions. Moreover, we thank Dr. Syuan-Ku Hsiao for his assistance with polymer synthesis, Dr. Krishna Gupta for his assistance with oligonucleotide purification, Isabell Jeglinski for synthesizing the modified RGD peptide, and Prof. Dr. Leoni Kunz-Schughart for providing U87MG_mcherry cells.

## Author contributions

E.K. conceived the project. Y.-H.P. and E.K. designed the experiments. Y.- H.P. carried out DNA sequence design, material characterization, and data analysis. Y.-H.P. and M.L. planned and carried out cell culture experiments. A.T. helped plan and interpret cell culture experiments. Y.-H.P. and E.K. wrote the initial paper draft. All authors discussed the results and helped revise the paper.

## Data availability

All data supporting the findings of this study are provided in the article and its Supplementary Information.

---

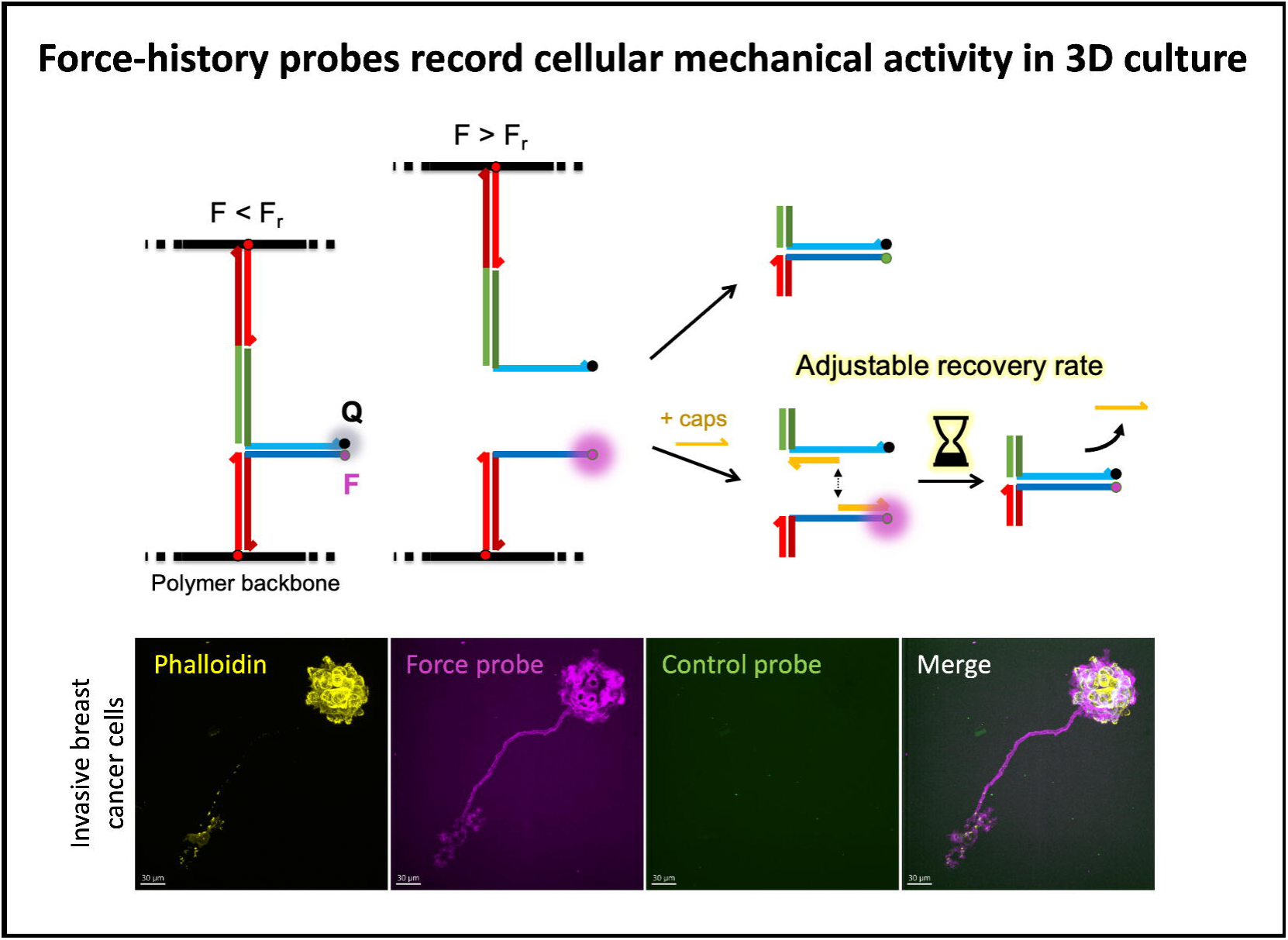

