## Supplementary Information for "Mapping 3D cellular mechanical activity with matrix-embedded DNA force-history probes"

### 1 Supplementary Notes

#### 1.1 Polymer mesh size calculation

To estimate the mesh size (ξ) of the polymer network, we applied the relationship $\xi \approx \nu_e^{-1/3}$, where $\nu_e$ is the effective crosslinker concentration in units of molecules per cubic meter, which is derived from the affine network model of rubber elasticity[1]. The following equation was used to calculate $\nu_e$ values. According to the affine network model, the elastic modulus, G', is directly proportional to the density of effective (i.e. inter-molecular) crosslinks, $\nu_e$[2]:

$$G' = A \cdot \nu_e \cdot R \cdot T$$

where R is the ideal gas constant, T is the absolute temperature in kelvin, and A is a constant that equals 1 for affine networks. The G' of hydrogel is ~ 135 Pa. The effective crosslinker concentration in hydrogels was estimated to be 52 µM ($\nu_e = 3.15 \times 10^{22}$ molecules/m³). Assuming a uniform, isotropic 3D network, the mesh size was calculated to be $\xi \approx (3.15 \times 10^{22})^{-1/3} \approx 32$ nm. This value represents the average distance between effective crosslinks within the molecular network for $\mathbf{P_{10}}$.

### 2 Supplementary Methods

#### 2.1 Solid-phase peptide synthesis (SPPS)

All SPPS chemicals were purchased from IRIS Biotech GmbH. RGD peptides (sequence: G(acryl-K)GGGRGDSP) were synthesized on a Liberty Blue HT12™ automatic and microwave-assisted peptide synthesizer (CEM GmbH) using a Rink Amide resin and a 9-fluorenylmethoxycarbonyl protection strategy. Amino acid activation was achieved by N, N'-diisopropyl carbodiimide, and ethyl cyanohydroxyiminoacetate. Acetylation of the N-terminus was performed by incubating the resin-bound peptides in acetic anhydride for 2 h. The reaction mixture was continuously stirred and vigorously saturated by bubbled nitrogen to prevent oxidation reactions and afterward washed three times with N, N-dimethylformamide. Deprotection of the amino acid side chains and cleavage from the resin was accomplished using a mixture of trifluoroacetic acid (ϕi = 87.5%), phenol (ϕi = 5%), triisopropylsilane (ϕi = 2.5%) and MilliQ-water (ϕi = 5%) for 3 h at room temperature. The crude peptide was precipitated in anhydrous diethyl ether, collected by vacuum filtration, and dried under nitrogen flow. Further purification of the peptides was realized by high-performance liquid chromatography (Agilent 1200, Agilent Technologies) on a preparative C18 column (10 µm particle size, 100 Å pore size, 250 x 30 mm, Phenomenex Ltd). A linear gradient of MilliQ-water/acetonitrile and trifluoroacetic acid (ϕi = 0.1%) was used as the mobile phase. Finally, the concentrated peptide-containing solution was lyophilized (Alpha 2-4 LD plus freeze-dryer, Martin Christ) and the obtained product was stored at -20 °C afterward until further usage.

#### 2.2 Polymer synthesis for additional cell culture

All experiments involving MDA-MB-231 cells and glioblastoma cells used a third derivative of the polymer $\mathbf{P_{20}^{RGD}}$, which contains higher degree of functionalization with anchor strands and lower polymer molecular weight. $\mathbf{P_{20}^{RGD}}$ contain the arginine-glycine-aspartate (RGD) peptide motif to facilitate cell adhesion. The synthesis protocol was adapted from refs. [3] and [4]. In brief, acrylamide (50 mg ml$^{-1}$), sodium acrylate (0.5 mg ml$^{-1}$), and acrylamide-labeled anchor strand DNA (Supplementary Table 1, strand 1) were co-polymerized at a molar ratio of 10,000:100:x (x = 20 for $\mathbf{P_{20}^{RGD}}$) in 1x TBE buffer (100 mM Tris, 90 mM boric acid, 1 mM EDTA, pH 8.3) to create DNA-grafted poly(acrylamide-co-acrylic acid). 10 mM acrylated RGD peptide (sequence: G(acryl-K)GGGRGDSP) was co-polymerized with the above solution. The synthesis of $\mathbf{P_{20}^{RGD}}$ was initiated by addition of 0.05 wt% TEMED and 0.05 wt% APS. To achieve high molecular weight and a narrow size distribution, the reaction was performed in high-purity nitrogen gas, which was passed through an oxygen trap on-site. The reaction was allowed to proceed overnight and yielded a highly viscous solution, indicating the formation of long polymer chains. NMR spectroscopy was used to verify high conversion of the monomer (Supplementary Figure 2). The solution was diluted in 9 volumes of 1x TE buffer (10 mM Tris, 1 mM EDTA, pH 8.0) and subsequently purified via methanol precipitation. The pellet was resuspended in milliQ water at 2.5% (w/v) and stored in aliquots at −20 °C.

#### 2.3 Nuclear magnetic resonance spectroscopy (NMR)

Nuclear magnetic resonance (NMR) spectroscopy was performed on samples of unpurified polymers prepared at a 1:8 dilution in $D_2O$. $^1$H NMR spectra were recorded at 30–32 °C on a 500 MHz spectrometer (Bruker) with a 2 second acquisition time and 32 transients. Chemical shifts (δ) are reported in parts per million (ppm) downfield from tetramethyl silane (TMS). The $^1$H NMR shifts are relative to the residual hydrogen peak of $D_2O$ (4.79 ppm).

#### 2.4 Rheological measurements

Viscoelasticity measurements were conducted using Anton Paar MCR301 rheometer with a 25-mm-diameter cone-plate geometry (cone angle 0.5 °). Temperature control was ensured by installing a Peltier system PTD-200 (Anton Paar) within the measurement area. To maintain high humidity and prevent evaporation artefacts, a wet paper cylinder was placed around the plate's circumference. All DyNAtrix samples were added onto the rheometer after freshly mixing the precursors. To initiate gelation, the temperature was first held at 4 °C for 5 min, then gradually increased to 20 °C over the course of 3 min, followed by a rise from 20 to 37 °C over 30 min. The temperature was then maintained at 37 °C for 240 min to complete gelation. All measurements were conducted at 37 °C. Frequency sweeps were conducted from 0.1 Hz to 100 Hz at 10% strain. Stress-relaxation properties were measured at 15% strain, where the strain was kept constant while recording shear stress over time (Supplementary Figure 3).

#### 2.5 2D culture of MDA-MB-231 cells

Breast cancer cells MDA-MB-231 were cultured in the serum-based media (Dulbecco's Modified Eagle Medium/Nutrient Mixture F-12 (DMEM/F-12) (Gibco, Cat. #11039047) supplemented with 10% Fetal Bovine Serum (FBS) (Sigma Aldrich, Cat. # F7524)) and 1x Penicillin and Streptomycin (Sigma Aldrich, Cat. #P4333) at 37 °C and 5% $CO_2$ in a humidified incubator. The cells were grown until 80%–90% confluence in T25 flask before detachment with 0.25% trypsin-EDTA solution. Collected cell suspension were washed and resuspended in serum-free medium at a density of 4 x $10^6$ cells/mL for embedding in DyNAtrix.

#### 2.6 Dye uptake test

Breast cancer cells MDA-MB-231 were seeded in a 96-well plate at 5000 cells/cm$^2$ and cultured in serum-free media (1x DMEM/F-12, 1x Insulin-Transferin-Selenium (Gibco, Cat. #41400045), 25 ng/mL EGF (Miltenyi, Cat. #130093825), 1x Penicillin and Streptomycin) for overnight. After cells have attached to the bottom, dye- and quencher-labeled DNA oligos were added to the serum-free media to reach 1 µM final concentration. Cells were incubated with DNA oligos for 4 days at 37 °C, 5% $CO_2$. Fluorescence images were acquired on Andor Dragonfly Confocal Microscope (Oxford Instrument) at 4x magnification (Supplementary Figure 10).

#### 2.7 Nuclease detection assay

The Förster-resonance-energy-transfer-paired oligos, modified with Cy5 fluorophore and Iowa Black Dark quencher (Q), were purchased from IDT. A mixture of 2 µM Cy5-strand (strand 20a) and 4 µM Q-strand (strand 20b) was prepared in 1x PBS buffer and subjected to annealing steps: heating at 95 °C for 1 min, instant cooling to 50 °C for 2 min and gradual cooling from 50 °C to 20 °C at a rate of −1.5 °C min−1. Cy5-Q paired oligos were added at 100nM final concentration in four different media conditions: (1) 10% FBS in 1x DMEM, (2) 1x DMEM, (3) the fresh serum-free medium (1x DMEM/F-12, 1x Insulin-Transferin-Selenium, 25 ng/mL EGF, 1x Penicillin and Streptomycin) and (4) the used serum-free medium obtained from 2D culture of MDA-MB-231 cells after two days. Each sample was loaded at a volume of 20 µL per well into a 96-well quantitative PCR plate. Fluorescence signals were recorded every 10 min for 24 hours at 37 °C using a Bio-Rad CFX96 Real-Time PCR System (Supplementary Figure 11).

#### 2.8 Preparation of DyNAtrix precursors for additional cell culture

For 3D culture of MDA-MB-231 cells, precursor A was prepared by mixing $\mathbf{P_{20}^{RGD}}$ (final concentration: 0.5% (w/v)), forward crosslinker (strand 2a, final concentration: 40 µM), blocking strand (strand 3a, final concentration: 80 µM), quencher-labeled probe strand (strand 5a, final concentration: 1 µM), fluorophore-labeled probe strand (strand 5b, final concentration: 2 µM). Precursor B was prepared by mixing reverse crosslinker (strand 2b, final concentration: 50 µM) and blocking strand (strand 3b, final concentration: 100 µM).

For 3D culture of glioblastoma spheroids, precursor A was prepared by mixing $\mathbf{P_{20}^{RGD}}$ (final concentration: 0.5% (w/v)), forward crosslinker (strand 2a, final concentration: 80 µM), blocking strand (strand 3a, final concentration: 160 µM), quencher-labeled probe strand (strand 5a, final concentration: 1 µM), fluorophore-labeled probe strand (strand 5b, final concentration: 2 µM). Precursor B was prepared by mixing reverse crosslinker (strand 2b, final concentration: 90 µM) and blocking strand (strand 3b, final concentration: 180 µM).

10x concentrated DMEM was added to both precursors to reach a final 1x concentration. All precursors were pre-annealed on a C1000 Touch™ Thermal Cycler (Bio-Rad) using the following steps: (1) heat to 70 °C for 3 min, (2) cool from 70 °C to 4 °C at −3 °C min$^{-1}$, (3) hold at 4 °C. The precursor was stored at 4 °C.

#### 2.9 3D culture of MDA-MB-231 cells

5 µL of the MDA-MB-231 cell suspension was first mixed with 15 µL precursor A on ice. 5µL precursor B was then added to the mixture (total volume 25 µL) and pipetted up and down for 25 times. The final cell density was 4 x $10^5$ cells/mL in DyNAtrix. The mixture was subjected to a micro-well insert (ibidi GmbH, Cat. #80409) placed in a 24-well plate with 5 µL cell-laden mixture per micro-well. The plate was incubated at 37 °C with 5% $CO_2$ for 60 min to trigger heat-activated gelation. Following gelation, 2 mL of serum-free media (1x DMEM/F-12, 1x Insulin-Transferin-Selenium, 25 ng/mL EGF, 1x Penicillin and Streptomycin) was added to each well. After overnight culture at 37 °C and 5% $CO_2$, the cells were stained with 1 µM CellTracker orange (Invitrogen, Cat. #C2927) in the serum-free medium for 30 min at 37 °C and 5% $CO_2$. Subsequently, the staining medium was removed and replaced with fresh serum-free medium containing 18-nt *split* DNA capping strands at a final concentration of 2 µM for F-strand caps (strand 18a), and 4 µM for Q-strand caps (strand 18b). Confocal images were acquired every 2 hours for 48 hours on an Andor Dragonfly Confocal Microscope (Oxford Instrument) at 20x magnification (Supplementary Video 4).

#### 2.10 Preparation for glioblastoma spheroids

U87MG_mcherry human glioblastoma cells were kindly provided by Prof. L. Kunz-Schughart, OncoRay—National Center for Radiation Research in Oncology, Dresden, Germany. Cells were maintained in T75 tissue culture flasks (BioLite Thermo Scientific) in Eagle's Minimum Essential Medium (Sigma-Aldrich, Cat. #M2279) supplemented with 10% Fetal bovine serum (Gibco, Cat. #A5256701) and 1x Penicillin and Streptomycin (Gibco, Cat. #15140122) at 37 °C, 5% $CO_2$ until 70–80% confluence before detachment with 1x TrypLE Express enzyme solution (Gibco, Cat. #11528856). Collected cell suspension were centrifuged and resuspended in culture medium at a density of $10^4$ cells/mL. To obtain spheroids composed of 1000 cells, 100 µl of this suspension were dispensed in each well of a Nuclon Sphera 96-well U-Shaped-Bottom Microplate (Thermo Scientific, Cat. #174929) followed by a quick centrifugation step (200 xg, 10s). After 72 hours of culture, spheroids were gently collected with a wide orifice pipet tip (Mettler Toledo) and embedded into DyNAtrix.

#### 2.11 3D culture of glioblastoma spheroids

5 µL of the glioblastoma spheroid suspension was first mixed with 15 µL precursor A on ice. 5 µL precursor B was then added to the mixture (total volume 25 µL) and pipetted up and down for 25 times. The final spheroid density was 200 spheroids/mL in DyNAtrix. The mixture was subjected to a micro-well insert (ibidi GmbH, Cat. #80409) placed in a 35 mm µ-dish (ibidi, Cat. #81158) with 5 µL cell-laden mixture per micro-well. The plate was incubated at 37 °C with 5% $CO_2$ for 60 min to trigger heat-activated gelation. Following gelation, 4 mL of serum-free media (1x DMEM/F-12, 1x Insulin-Transferin-Selenium, 1x Penicillin and Streptomycin) was added. After 4 hours incubation at 37 °C and 5% $CO_2$, 18-nt *split* DNA capping strands were added to the medium at a final concentration of 2 µM for F-strand caps (strand 18a), and 4 µM for Q-strand caps (strand 18b). Confocal images were acquired every 30 minutes for 24 hours on an Andor Dragonfly Confocal Microscope (Oxford Instrument) at 20x magnification (Supplementary Video 5).

#### 2.12 DNA rupture force calculations

We employed a recently developed tool from Liu and Yan[5,6] to estimate the approximate force at which the force-history probe would get activated. The model was calibrated under physiological ionic strength (150 mM NaCl), pH (7.4), and temperature (37 °C), which matches the experimental conditions in mammalian cell culture. We set the loading rate to 1 pN/s, which lies within the relevant range of loading rates experienced by integrins[7]. An ideal zipper opening geometry was used for assessing the rupture force of the force probe domain (CGTTCAAAGATGTTTCAAATTCAACG), whilst an ideal shear opening geometry was used for assessing the rupture force of the anchor (GACGGCTCATAAGGCTCTAATC) and overlap domain (TGTGTTAGTCANTGTCCCATTA, where N was replaced by an explicit A nucleotide). The obtained rupture forces are 3.3 pN for the force probe, 30 pN for the anchor, and 26 pN for the overlap domain (Supplementary Figure 4).

#### 2.13 Dimer formation propensity calculations

We estimated the propensity of cap dimer formation using the thermodynamic prediction tool NUPACK[8]. The predictions were run under physiologically relevant parameters, with the salt concentration of 150 mM NaCl and at the temperature of 37 °C. The in silico mixture consisted of equimolar concentrations (1 µM) of the forward and reverse capping strands. Based on the predicted equilibrium concentrations of the assembled complexes, the propensity of dimerization was calculated as the ratio of the dimer concentration to the total DNA concentration (Supplementary Figure 14).

#### 3 Supplementary Figures

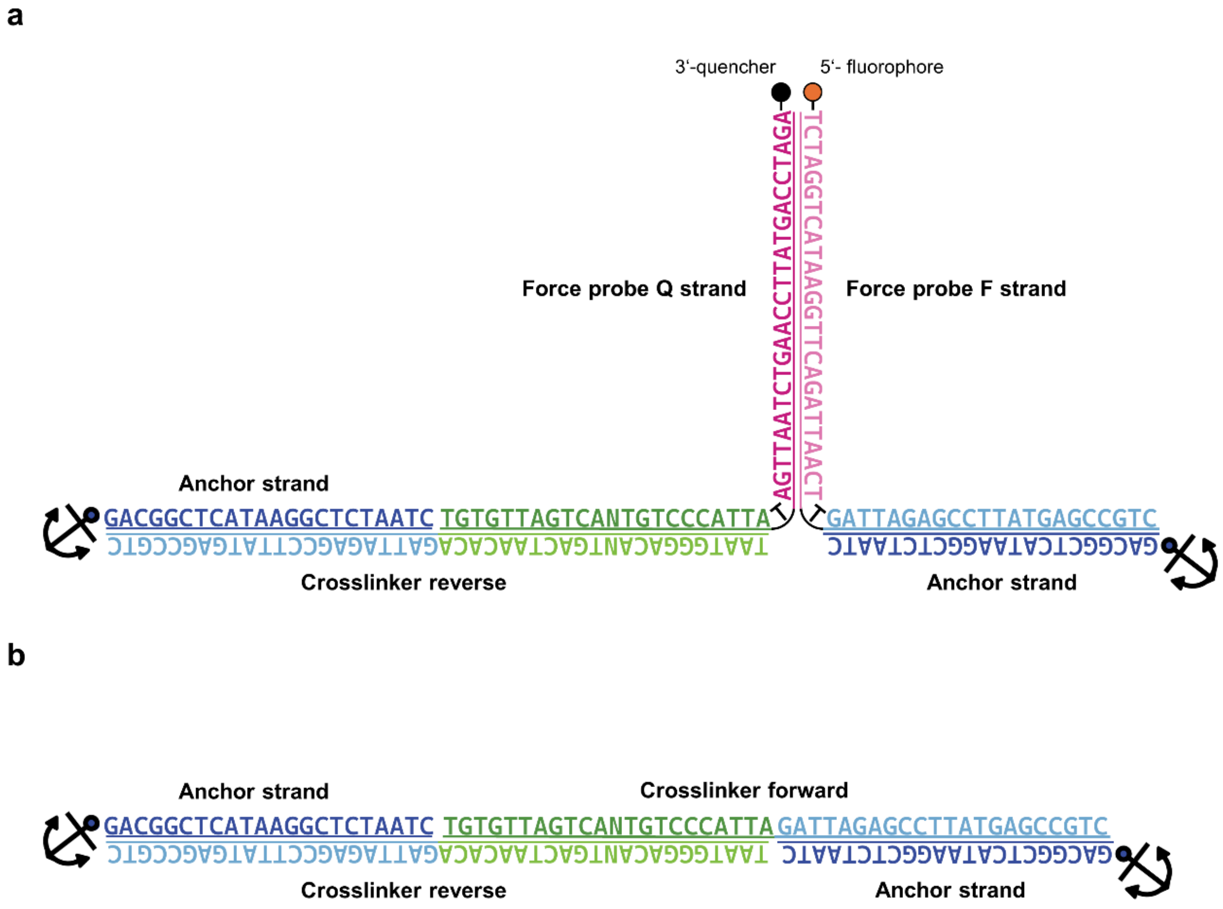

**Supplementary Figure 1. Schematic representation of DNA crosslinks in DyNAtrix. a)** Force-responsive crosslinks contain a 26-nt probe domain (magenta), a 22-nt crosslinker overlap domain (green), and an adaptor domain (light blue) that hybridize to the anchor strand (dark blue) conjugated on the polymer. The use of ambiguous bases (N) diversifies the overlap domain and increases the efficiency of intermolecular crosslinking[4,9]. The probe domain is modified with a fluorophore–quencher pair at the termini that separates upon mechanical unzipping[10,11]. **b)** Dual-splinted DNA crosslinks with a 22-nt overlap domain. Anchor symbols indicate the position at which the anchor strands are covalently bound to the poly(acrylamide-co-acrylic acid) backbone.

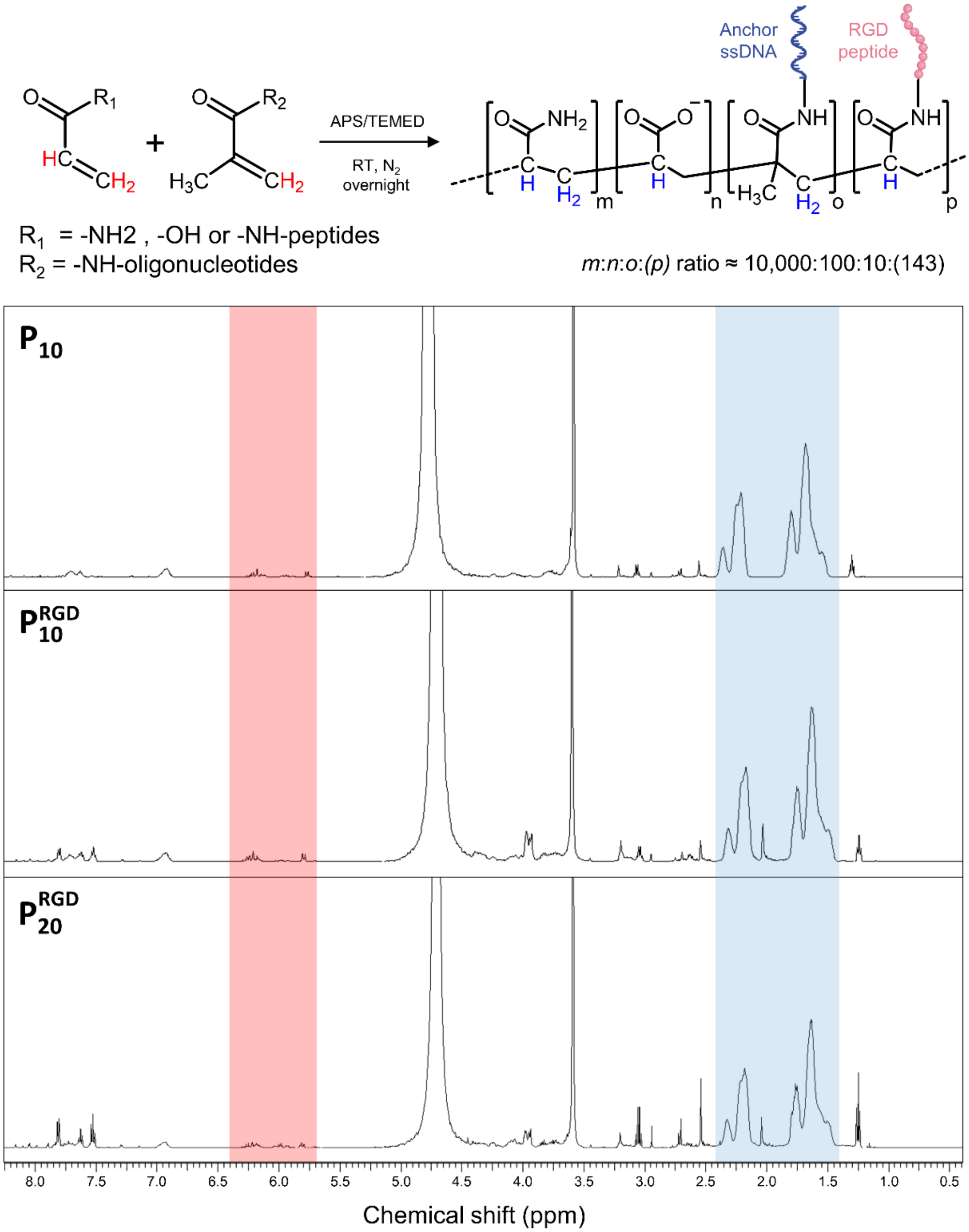

**Supplementary Figure 2. $^1$H-NMR spectra of $P_{10}$, $P_{10}^{RGD}$ and $P_{20}^{RGD}$ in $D_2O$ prior to methanol purification.** The conversion percentage is determined by measuring the ratio of free residual acrylamide monomer protons (δ ~5.7–6.4; orange) to polymer backbone protons (δ ~1.4–2.4; blue). **$P_{10}$** showed 98.4% conversion; **$P_{10}^{RGD}$** showed 98.8% conversion; **$P_{20}^{RGD}$** showed 98.9% conversion.

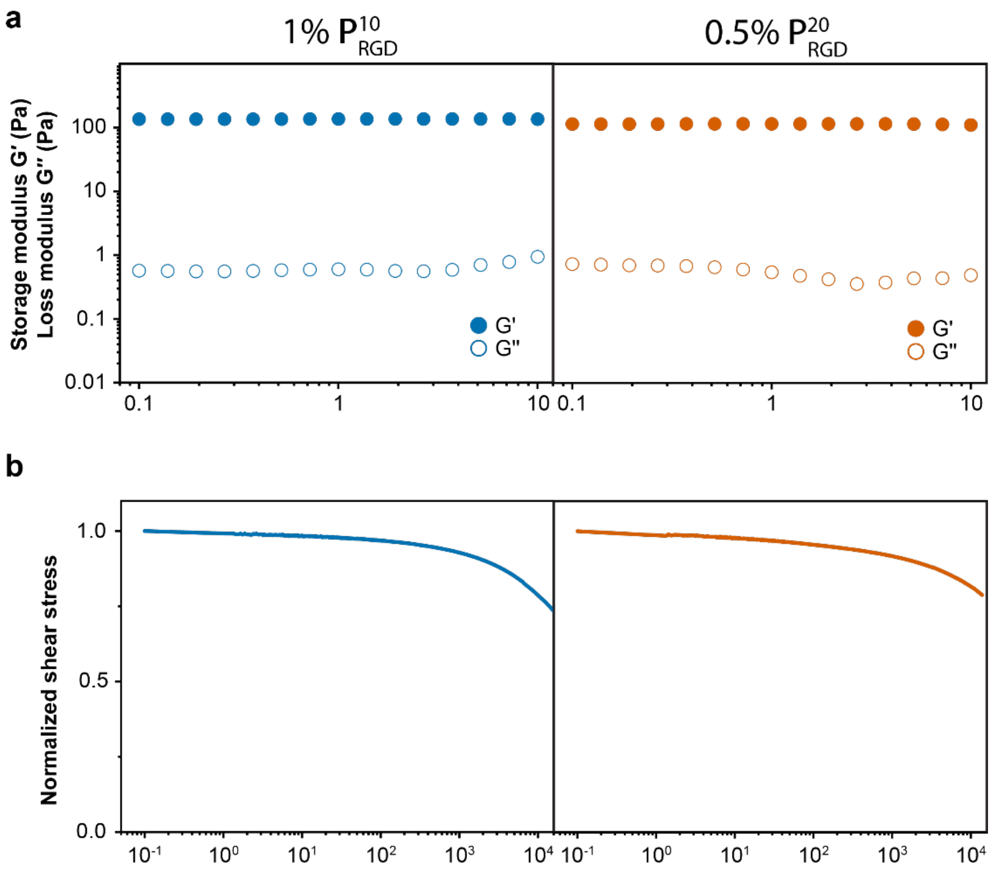

**Supplementary Figure 3. Rheological characterization of the hydrogels used in cell culture experiments. a)** Frequency sweeps and **b)** normalized stress-relaxation curves of DyNAtrix prepared with 1% (w/v) $\mathbf{P_{10}^{RGD}}$ and 0.5% (w/v) $\mathbf{P_{20}^{RGD}}$ crosslinked with 80 µM crosslinker library (CCL-4, 22nt). The former was used in MCF10A-Er-Src cell culture, the latter was used in glioblastoma cell culture. All measurements were carried out at 37 °C. Shear moduli and stress relaxation are governed by the concentration and nanomechanical stability of the crosslinkers[4,12].

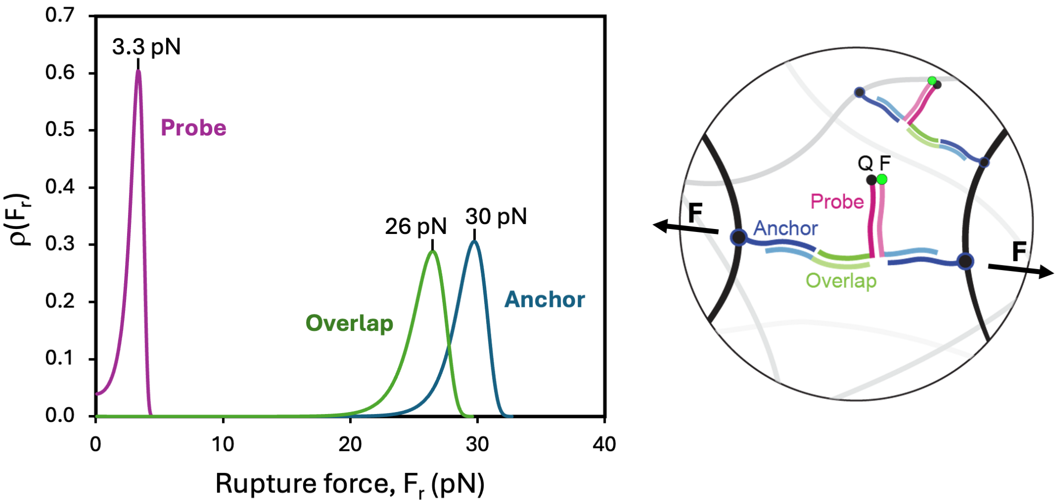

**Supplementary Figure 4. Probability density functions of the predicted rupture forces of different FHP domains at 37 °C and 1 pN/s loading rate.** The FHP can, in principle, rupture in 3 different positions. The probability density function, ($F_r$), was calculated for each of the three domains separately. Expectedly, the probe domain is predicted to break at 3.3 pN, well before the breakage of overlap domain breakage (26 pN) and anchor breakage (30 pN) would occur.

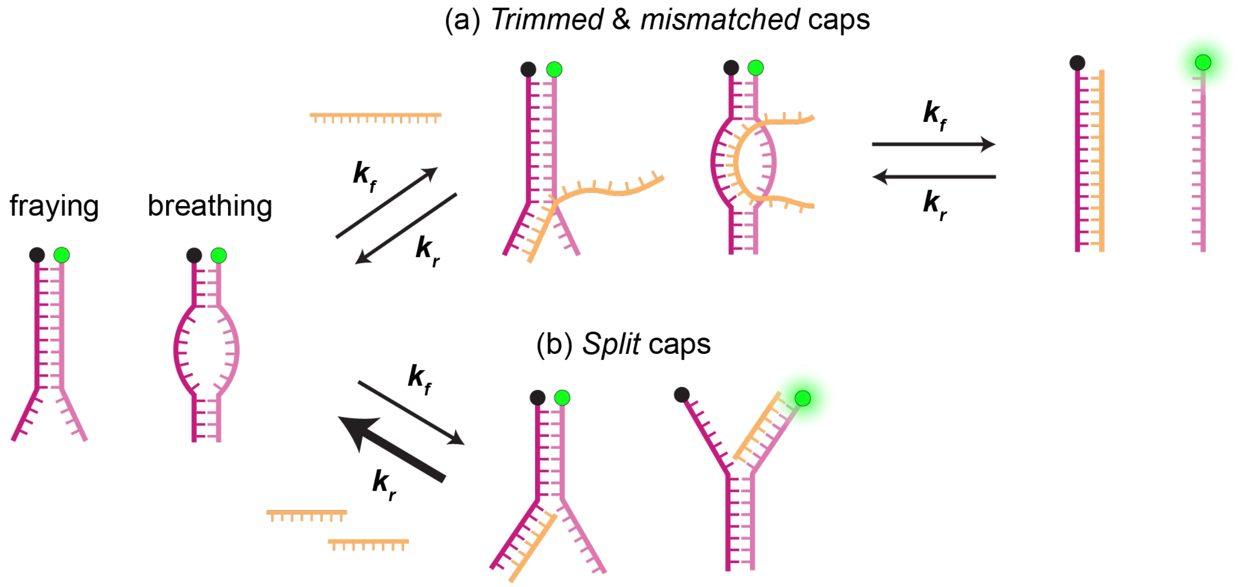

**Supplementary Figure 5. Background fluorescence caused by spurious "leak" reactions. a)** Leakage is initiated by end-fraying and breathing of the probe duplex, which exposes bases for cap binding and subsequent strand displacement. Over time, the forward reaction (cap invasion) and the reverse reaction (probe re-hybridization) reach an equilibrium, yielding a constant fraction of opened probes and a corresponding background fluorescence plateau. **b)** Though spurious binding of the *split* caps to the probe is possible, their reduced length does not readily cause full dissociation of the probe.

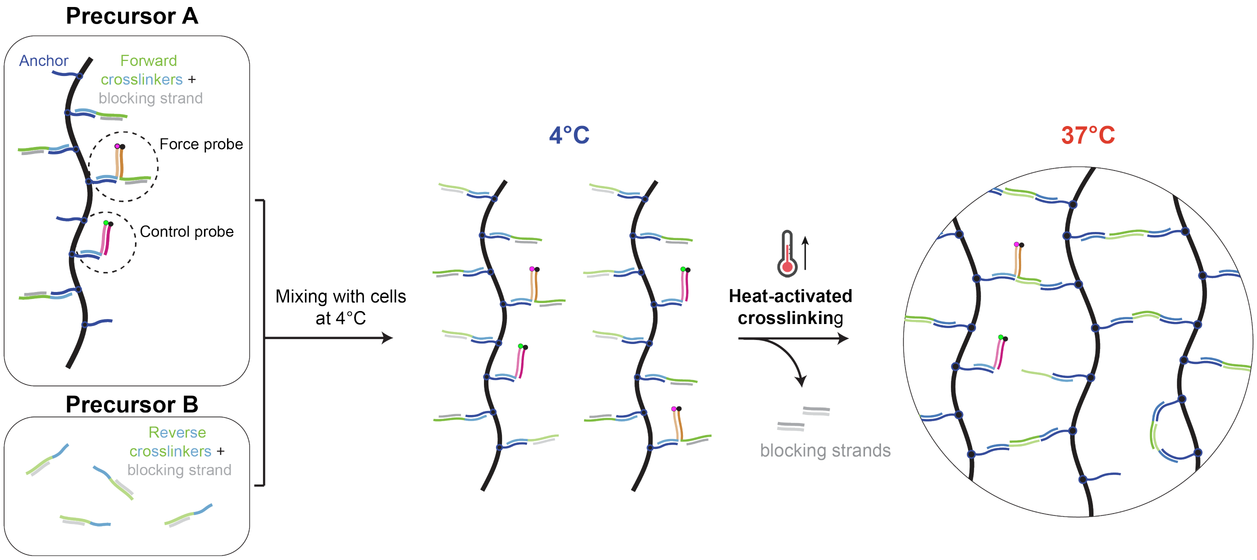

**Supplementary Figure 6. Heat-activated gelation procedure for cell encapsulation.** Two liquid precursors A and B are prepared. Precursor A contains the 1% (w/v) DNA-functionalized polymer $\mathbf{P_{10}^{RGD}}$ or $\mathbf{P_{20}^{RGD}}$, the forward crosslinker, the blocking strands, and 1 µM force probe and control modules; B contains the reverse crosslinker and the corresponding blocking strands. Both precursors were pre-annealed to ensure complete binding of blocking strands to the crosslinkers and were subsequently stored at 4 °C until use. Upon mixing two precursors with a cell suspension and warming to 37 °C, the blocking strands dissociate, triggering crosslinking and rapid gelation. Heat-activated crosslinking enables gentle cell encapsulation within a homogenous matrix using a protocol analogous to Matrigel handling.

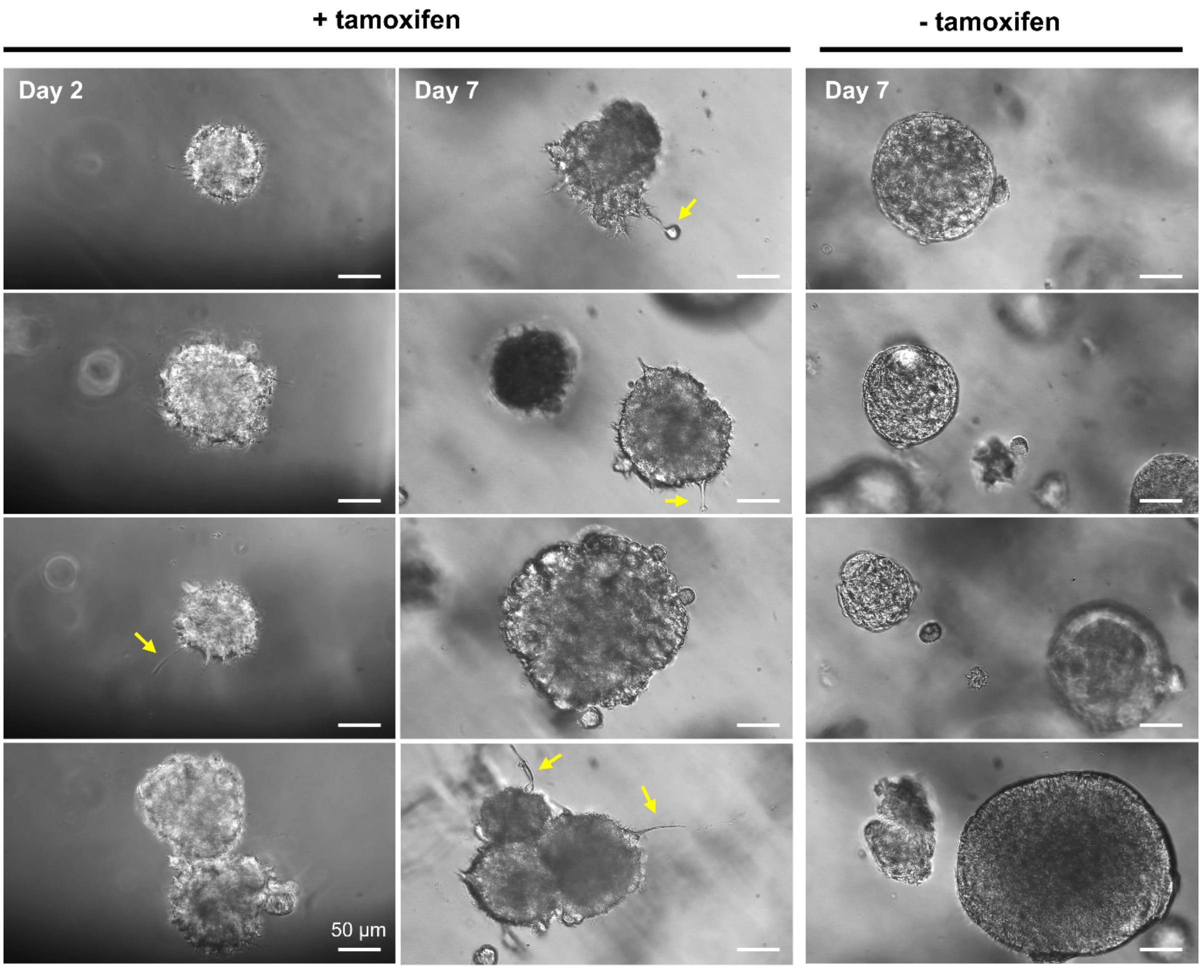

**Supplementary Figure 7. Tamoxifen supplementation induces an elongating, invasive phenotype in MCF-10A ER-Src cells.** Brightfield images of breast cancer spheroids MCF-10A ER-Src cultured in DyNAtrix in the presence or absence of 1 µM tamoxifen in the serum-free medium. Tamoxifen-treated cells display long protrusion arms (yellow arrows) and localized single-cell invasion into the matrix (top middle). In contrast, the untreated spheroids remained compact and rounded. Scale bar = 50 µm.

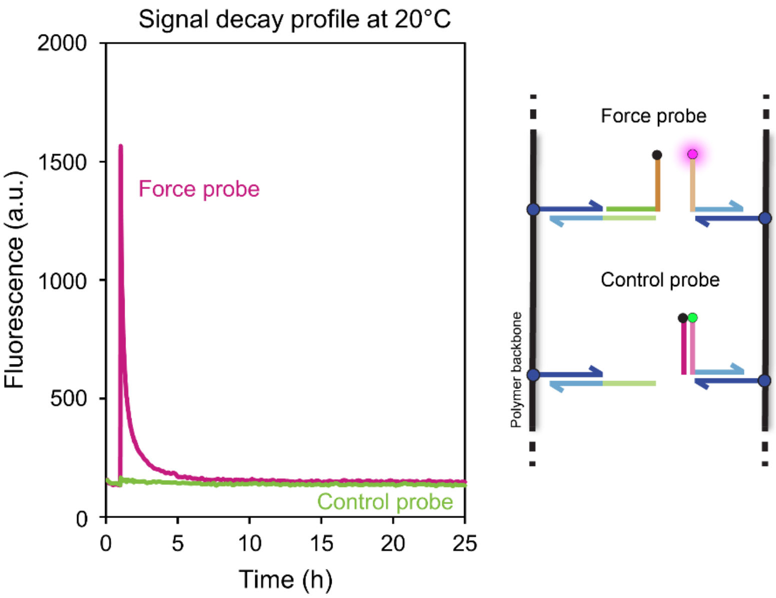

**Supplementary Figure 8. Validation of force probe vs. control**. Fluorescence profiles of the functional force probe 1 (strands 4a and 4b) and control probe (strands 4c and 4b) following mechanical perturbation. The control probe, which lacks the crosslinker's overlap domain necessary to bridge the polymer network, shows no activation signal, confirming that the functional probe signals are driven by mechanical tension transmitted through the crosslinks.

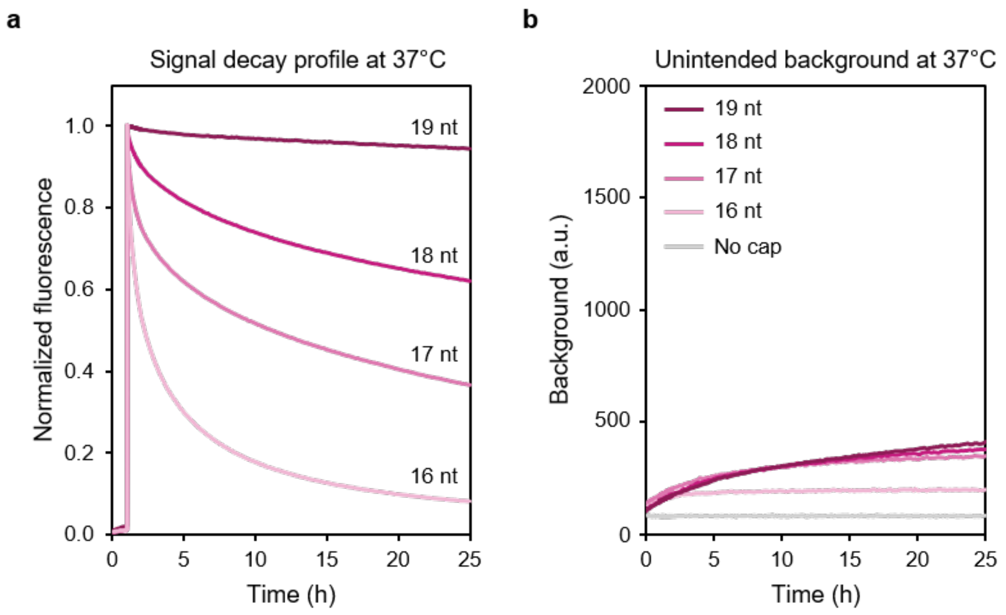

**Supplementary Figure 9. Tuning of signal decay rate with the *split* capping strands at physiological temperature. a)** Normalized fluorescence decay profiles of force probe 2 in the presence of *split* capping strands (16–19 nt) at 37 °C. **b)** background fluorescence accumulation over 25 hours. The data shows that while longer capping strands extend signal retention, they also mildly increase the baseline background due to spontaneous thermal breathing at 37 °C.

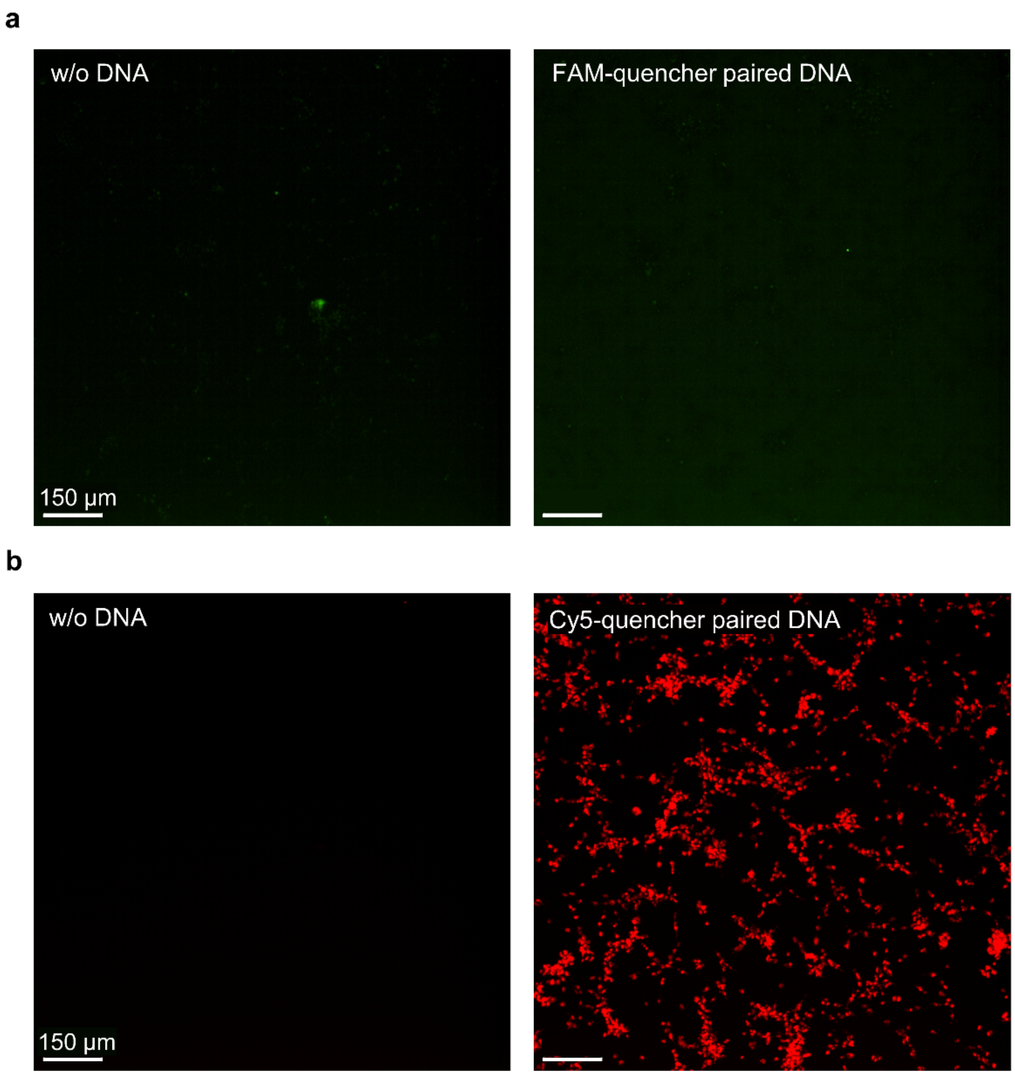

**Supplementary Figure 10. Evaluation of fluorophore uptake by breast cancer cells.** Confocal images of MDA-MB-231 cells cultured on a 2D substrate and incubated with 1 µM of DNA oligos modified with **a)** quencher-labeled strand (strand 4a) and FAM-labeled strand (strand 4b) or **b)** quencher-labeled strand (strand 20b) and Cy5-labeled strand (strand 20a) for 4 days in serum-free media. The control group was incubated without DNA oligos and imaged in the FAM and Cy5 channels. The cell bodies exhibited strong signal in the presence of the Cy5-quencher FRET-paired oligos, indicating a significant uptake of Cy5 molecules by the cells. Scale bar = 150 µm.

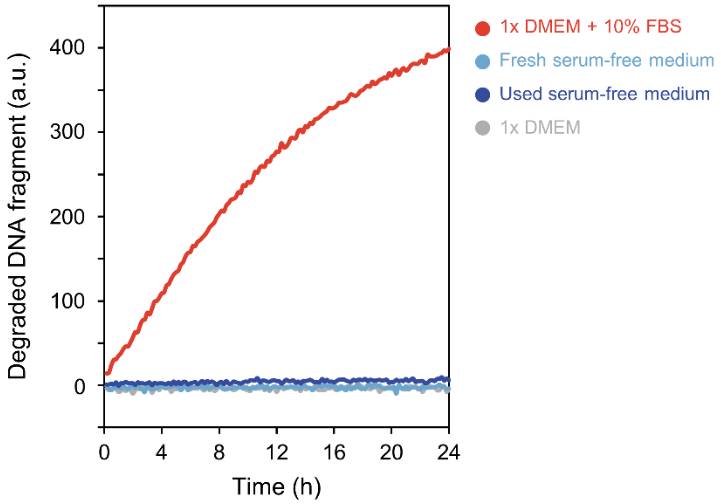

**Supplementary Figure 11. Stability of DNA fluorescent probes in cell culture media.** Nuclease detection assay using the increase in force probe fluorescence (strands 20a and 20b) to quantify the rate of probe degradation in FBS-supplemented vs. serum-free media over 24 hours at 37 °C. To quantify cell-secreted nucleases, the used serum-free medium was obtained after 2 days of 2D culture with MDA-MB-231 cells. Minimal degradation is observed in serum-free media with and without prior incubation with the cells, validating the stability of the probes for multi-day cell culture experiments.

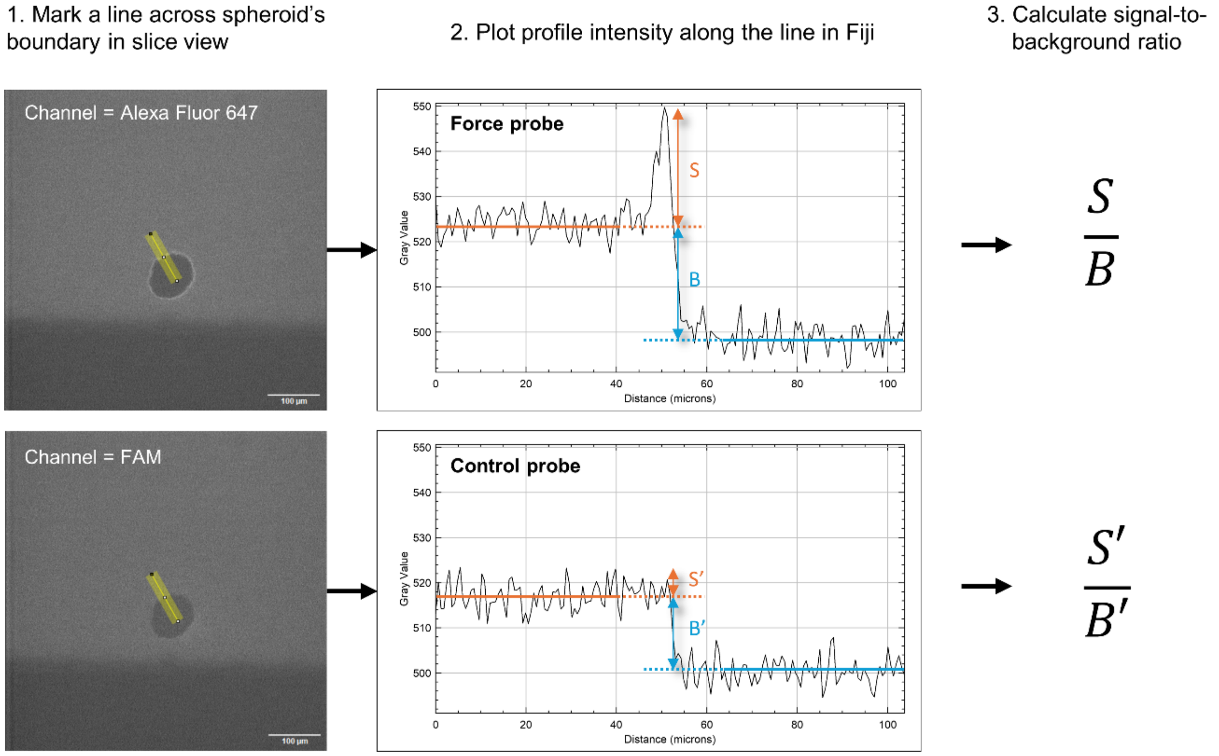

**Supplementary Figure 12. Quantification of signal-to-background (S/B) ratios around spheroids.** Intensity profiles are plotted in Fiji[13] across the spheroid–gel interface in the confocal sections. The same ROI is quantified in both force probe and control channels. The signal (S) is defined as the peak intensity at the boundary relative to the bulk gel intensity extrapolated from the gel region (the first 40 µm of the curve). The background (B) was calculated as the difference between the extrapolated gel intensity and the baseline intensity extrapolated from the dark lumen regions (the last 40 µm of the curve).

a

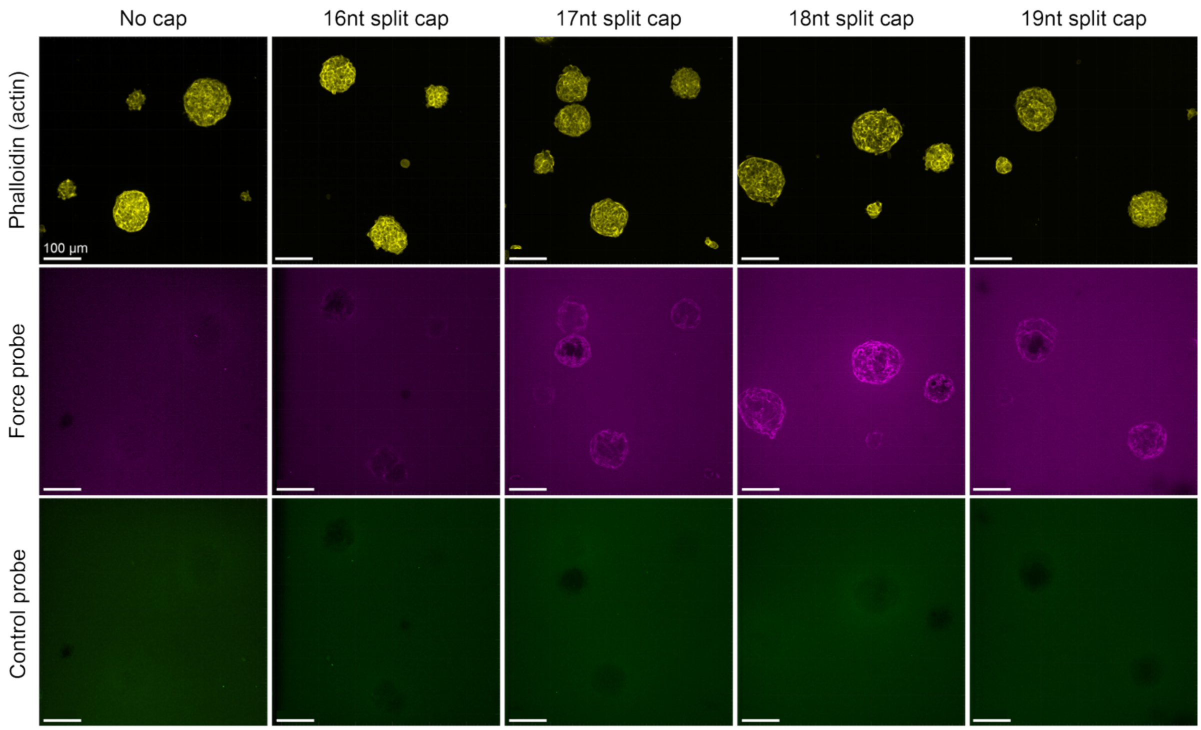

b

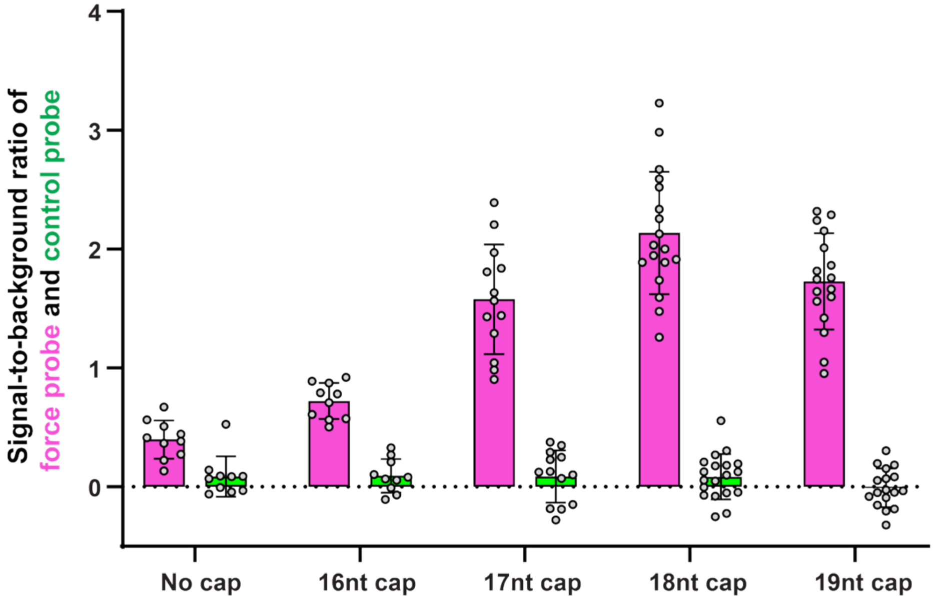

**Supplementary Figure 13. Comparison of probe signal activated by spheroid expansion. a)** Representative confocal images of MCF-10A ER-Src spheroids cultured with force probes in the presence of varying lengths of *split* capping strands (16–19 nt). Scale bar = 100 µm. **b)** Quantification of S/B ratios. 18-nt capping strands yield the strongest signal, whereas the control signal remains low across all samples.

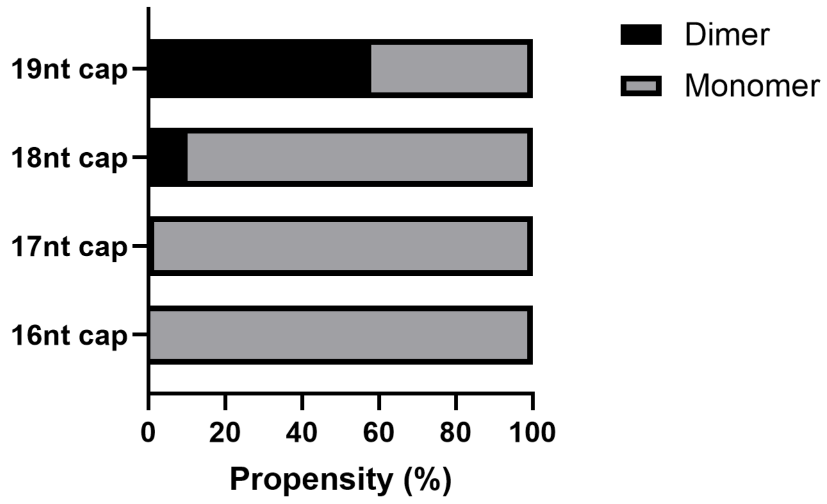

**Supplementary Figure 14. Comparison of dimer formation propensity between 16–19nt *split* caps.** 19nt caps are more prone to dimerize (58%) than 18nt caps (10%), 17nt caps (1.5%), and 16nt (<1%). The dimerization of caps is undesired, as it reduces their ability to rapidly bind activated probes. The analysis was carried out in NUPACK[8].

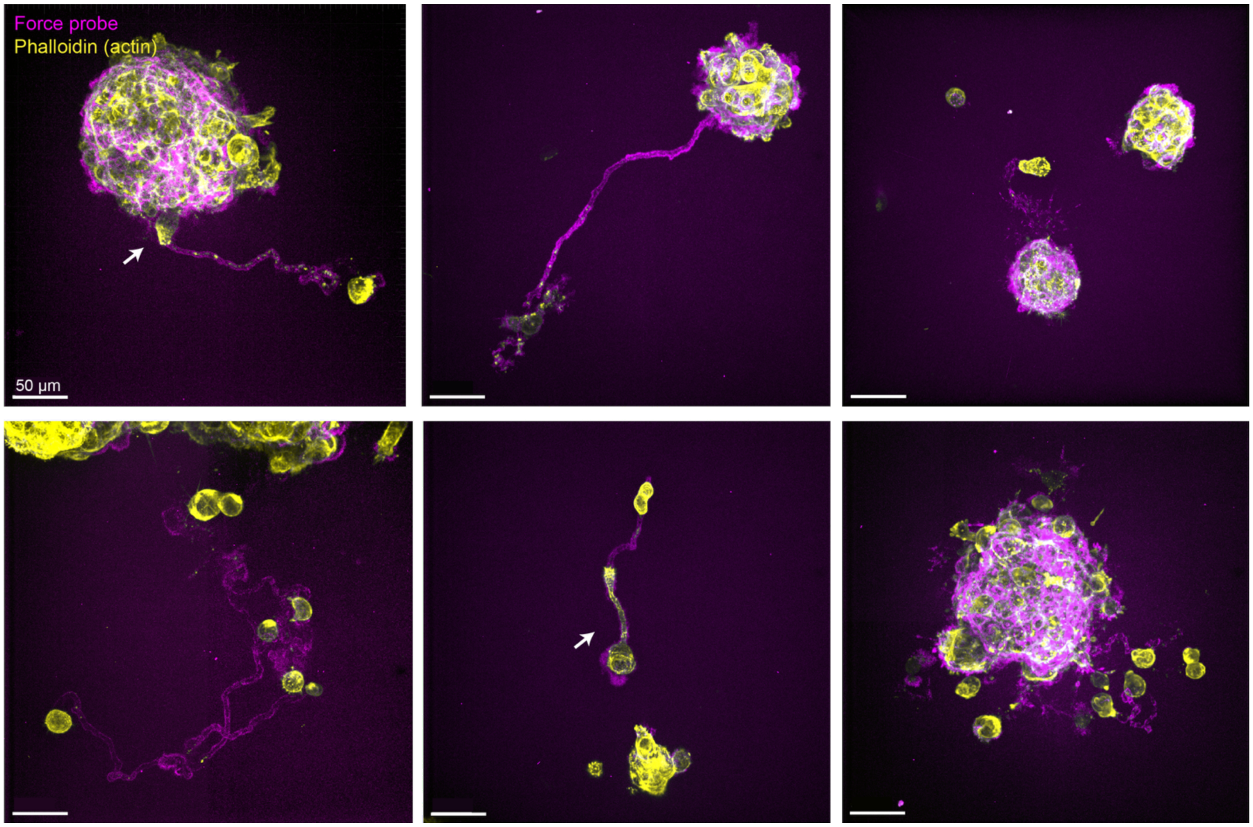

**Supplementary Figure 15. Confocal images of diverse fluorescent tracks activated by migrating cells.** Breast cancer spheroids MCF10AER-Src were cultured in DyNAtrix with force probes in the presence of 18-nt *split* capping strands for 6 days and subsequently fixed and stained with Alexa Fluor 555 Phalloidin (yellow). The activation fluorescence (magenta) marks the diverse trajectories of the migrating cells. In some instances, migrating cells were observed to precisely follow the trajectories of earlier cell migrations (white arrows). Scale bar = 50 µm.

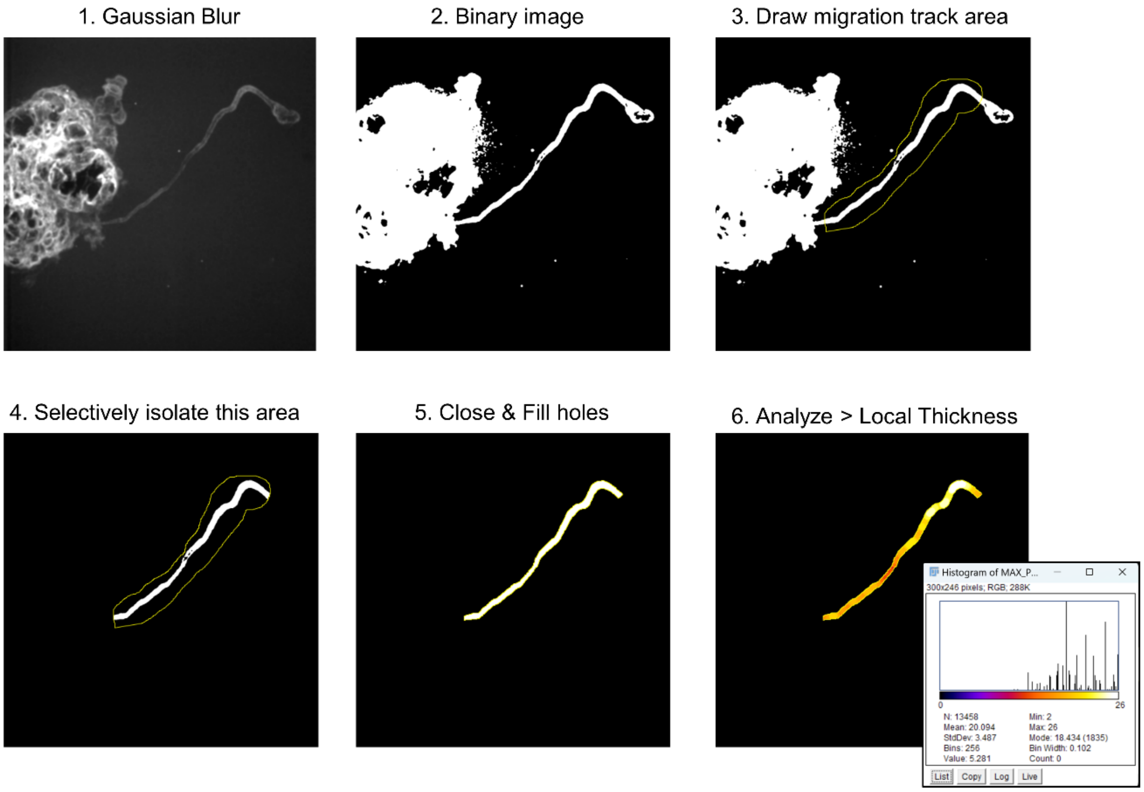

**Supplementary Figure 16. Thickness analysis of cell migration channels.** Maximum projection of confocal images containing cell-migration tracks were analyzed in Fiji[13]. The mean thickness of migration tracks was measured using the Local Thickness function. The migration channels displayed relatively uniform diameters of 6.02 ± 0.26 µm. Data is shown as mean ± s.d. (n = 3 migrating cells).

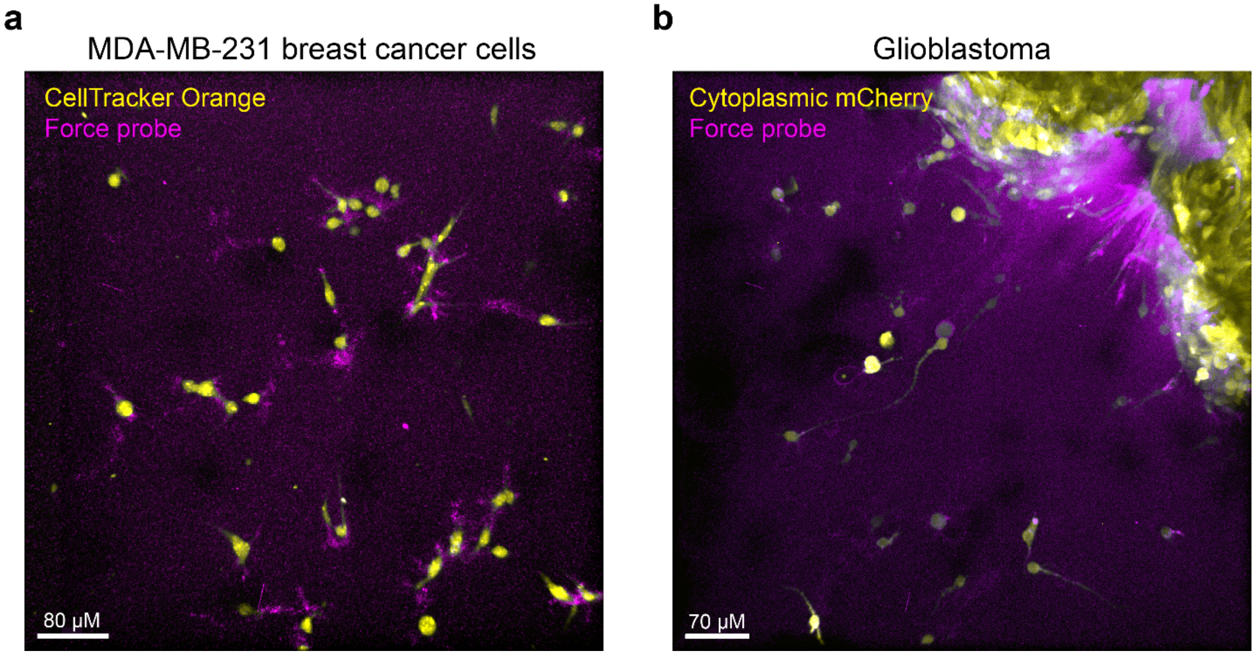

**Supplementary Figure 17. Force-history tracking of glioblastoma and breast cancer cell models.** **a)** MDA-MB-231 breast cancer cells were embedded in DyNAtrix and cultured in serum-free medium with 18-nt *split* capping strands. Cells were stained with CellTracker orange (cytoplasm). **b)** Glioblastoma spheroids (U87MG_mcherry cells) were embedded in DyNAtrix and cultured in serum-free medium with 18-nt *split* capping strands. Cells at the spheroid periphery actively protruded and left long, spindle-like marks of probe activation (upper right corner). Some cells invaded the matrix, generating variable shapes and sizes of probe activation marks.

#### 4 Supplementary Tables

**Supplementary Table 1. List of DNA sequences used in this study.** Blue: anchor strand sequence or adaptor site complementary to the anchor strand; green: crosslinker overlap domain; ochre yellow: ambiguous N base (A, T, C, or G); magenta: force probe domain; red: mismatched base. 5Acryd: Acrydite. 3IABkFQ: Iowa Black® quencher FQ; 56-FAM: Fluorescein; 3IAbRQSp: Iowa Black® quencher RQ; 5Alex647N: Alexa Fluor® 647.

| Strand | Anchor strand | nt |
|---|---|---|
| 1 | /5Acryd/GACGGCTCATAAGGCTCTAATC | 22 |
| | **Crosslinker and its blocking strand for heat-activated gelation** | |
| 2a | TGTGTTAGTCANTGTCCCATTAGATTAGAGCCTTATGAGCCGTC | 44 |
| 2b | TAATGGGACANTGACTAACACAGATTAGAGCCTTATGAGCCGTC | 44 |
| 3a | ACATGGGACANAGACTAACATT | 22 |
| 3b | AATGTTAGTCTNTGTCCCATGT | 22 |
| | **Force probe 1 (FAM channel)** | |
| 4a | TGTGTTAGTCANTGTCCCATTATCGTTCAAAGATGTTTCAAATTCAACG/3IABkFQ/ | 49 |
| 4b | /56-FAM/CGTTGAATTTGAAACATCTTTGAACGTGATTAGAGCCTTATGAGCCGTC | 49 |
| | **Truncated sequence of force probe 1 as control** | |
| 4c | CGTTCAAAGATGTTTCAAATTCAACG/3IABkFQ/ | 26 |
| | **Force probe 2 (Alexa Fluor 647 channel)** | |
| 5a | TGTGTTAGTCANTGTCCCATTATAGTTAATCTGAACCTTATGACCTAGA/3IAbRQSp/ | 49 |
| 5b | /5Alex647N/TCTAGGTCATAAGGTTCAGATTAACTTGATTAGAGCCTTATGAGCCGTC | 49 |
| | **Trimmed capping strands for force probe 1** | |
| 6 | CGTTGAATTTGAAACATCTTTGAACG | 26 |
| 7 | GTTGAATTTGAAACATCTTTGAAC | 24 |
| 8 | TTGAATTTGAAACATCTTTGAA | 22 |
| | **Mismatched capping strands for force probe 1** | |
| 9 | CGTTGAATTTGATACATCTTTGAACG | 26 |
| 10 | CGTTGAATTAGAAACAACTTTGAACG | 26 |
| | **Split capping strands for force probe 1** | |
| 11a | CGTTGAATTTGAAACA | 16 |
| 11b | CGTTCAAAGATGTTTC | 16 |
| | **Split locking capping for force probe 2** | |
| 12a | CTAGGTCATAAG | 12 |
| 12b | GTTAATCTGAAC | 12 |
| 13a | TCTAGGTCATAAG | 13 |
| 13b | AGTTAATCTGAAC | 13 |
| 14a | TCTAGGTCATAAGG | 14 |
| 14b | AGTTAATCTGAACC | 14 |
| 15a | TCTAGGTCATAAGGT | 15 |

| | | |
|---|---|---|
| 15b | AGTTAATCTGAACCT | 15 |
| 16a | TCTAGGTCATAAGGTT | 16 |
| 16b | AGTTAATCTGAACCTT | 16 |
| 17a | TCTAGGTCATAAGGTTC | 17 |
| 17b | AGTTAATCTGAACCTTA | 17 |
| 18a | TCTAGGTCATAAGGTTCA | 18 |
| 18b | AGTTAATCTGAACCTTAT | 18 |
| 19a | TCTAGGTCATAAGGTTCAG | 19 |
| 19b | AGTTAATCTGAACCTTATG | 19 |
| | **Nuclease digestion probe** | |
| 20a | /5Cy5/CCGAGGACTGAGGGTTTTAGGAGTTGGTCTATAATCATGG | 41 |
| 20b | AGACCAACTCCTAAAACCCTCAGTCCTCGG/3IAbRQSp/ | 30 |
